# Cryo-ET of a human GBP coatomer governing cell-autonomous innate immunity to infection

**DOI:** 10.1101/2021.08.26.457804

**Authors:** Shiwei Zhu, Clinton J. Bradfield, Agnieszka Mamińska, Eui-Soon Park, Bae-Hoon Kim, Pradeep Kumar, Shuai Huang, Yongdeng Zhang, Joerg Bewersdorf, John D. MacMicking

## Abstract

All living organisms deploy cell-autonomous defenses to combat infection. In plants and animals, these activities generate large supramolecular complexes that recruit immune proteins for protection. Here, we solve the native structure of a massive antimicrobial complex generated by polymerization of 30,000 human guanylate-binding proteins (GBPs) over the entire surface of virulent bacteria. Construction of this giant nanomachine takes ∼1-3 minutes, remains stable for hours, and acts as a cytokine and cell death signaling platform atop the coated bacterium. Cryo-ET of this “coatomer” revealed thousands of human GBP1 molecules undergo ∼260 Å insertion into the bacterial outer membrane, triggering lipopolysaccharide release that activates co-assembled caspase-4. Together, our results provide a quasi-atomic view of how the GBP coatomer mobilizes cytosolic immunity to combat infection in humans.

**One-Sentence Summary:** Thousands of GBPs coat cytosolic bacteria to engineer an antimicrobial signaling platform inside human cells.

## Main Text

In biological systems, environmental cues are often sensed by ligand-induced allosteric changes in cell surface receptors that rapidly transmit signals to the interior for mobilizing the desired physiological response. Within the cell-autonomous defense systems of plants and animals (1, 2), additional modalities are used to decode the outside world including higher-order protein assemblies formed via helical symmetry (3–5). These intracellular assemblies amplify signals through protein polymerization or “prionization” events to facilitate proximity-induced autoactivation of latent zymogens, caspases and kinases. The benefits to such repetitive design are manifold: lowered signaling thresholds, all-or-none responsivity, and stable signalosome platforms that can recruit and accommodate numerous protein partners (4).

This ability to generate extended, filamentous signaling platforms stems in part from the modularity of the proteins involved. Signalosome proteins often harbor leucine-rich repeat (LRR) domains, caspase-activation and recruitment domains (CARDs), and death effector domains (DEDs) that concentrate receptors, adaptors and effectors through co-operativity (4). As a result, they yield some of the most iconic and visible structures inside immune-activated cells. These include RIG-I filaments and MAVs prion-like structures that control RNA sensing; ASC and NLRC4 filaments as part of the inflammasome machinery; Myddosomes orchestrating NF-*κ*B and IRF responses; and NLR ZAR pentamers that underpin the plant resistosome (3, 6). Collectively these large polymeric structures represent an increasingly pervasive paradigm for cell-autonomous immunity throughout metazoan evolution. Their mode of assembly differs from classical antigen or immunoglobulin signaling at the plasma membrane that are typically transmitted via clustered synapses (4).

Here we characterize a massive antimicrobial defense complex comprising over 30,000 proteins at the cryo-electron tomography (cryo-ET) level. Eukaryotic immune GTPases termed guanylate-binding proteins (GBP) assemble large polymeric structures inside *Arabidopsis*, zebrafish, mouse and human cells, in some cases using phase-separation to further concentrate homotypic complexes (7–11). In plants, GBP-like proteins (GBPLs) respond to inducible immune signals including salicylic acid and pipecolic acid to assemble large ∼200-400nm nuclear RNA polymerase II hubs that transcribe host defense genes following infection (7). In animals, immune cytokines such as interferons (IFNs) induce GBP expression to control microbicidal or inflammasome responses within the host cell cytosol (8,9,12); these activities often require relocation of GBPs to the site of microbial replication where they completely “coat” targeted pathogens to build mesoscale signaling or killing platforms (8,11–15). Such GBP-coated pathogens can range in size from ∼750nm in diameter for *Salmonella enterica* serovar Typhimurium (*Stm*) to >10μm for *Toxoplasma gondii* (16).

Remarkably, despite their central role for host defense across plant and animal kingdoms (7, 17), the ultrastructural and functional organization of these mesoscale GBP coatomers is unknown. Visualizing GBP complexes below the light diffraction limit would therefore enhance our understanding of how eukaryotic cells recognize and combat infection. Elucidating GBP assembly on the pathogen surface under physiological conditions and whether pathogens sense the presence of this massive coat are additional questions currently unanswered. Each have implications for anti-infective therapy within the human population as well as the basic biology of innate immune recognition.

### Human GBP1 coatomer size, kinetics, and stability *in situ*

We first examined the human GBP coatomer surrounding cytosolic bacteria at sub-diffraction limits using whole-cell 4Pi single-molecule switching nanoscopy (W-4PiSMSN) and fast, live three-dimensional (3D) OMX-SR imaging equipped with high-speed galvanometers. W-4PiSMSN is a dual objective iPALM/4Pi-SMSN nanoscope resolving 3D structures to ∼5-10nm (50-100 Å) linearly and axially throughout entire mammalian cells (18); it enabled us to detect single GBP molecules on the surface of virulent *Stm* inside human cells. 3D-Blaze super-resolution microscopy and wide-field imaging is capable of ∼180 and >300 frames.sec^-1^, respectively, ensuring coatomer assembly could be followed throughout the entire bacterial encapsulation process.

We tracked GBP1 as the forerunner of this 7-member dynamin-like GTPase family in humans. Recent work discovered GBP1 recruitment onto cytosol-invasive pathogens including *Stm* and *Shigella flexneri* enables bactericidal activity by apolipoprotein L3 in IFN-*γ*-activated primary human intestinal epithelia, myofibroblasts and endothelium, as well as in HeLa CCL2 cells (12). We and others also reported that GBP1 recruits additional GBP family members plus endogenous human caspase-4 to stimulate cytokine release (interleukin-18; IL-18) and pyroptosis in primary intestinal human organoids, human macrophages, and human epithelial cell lines (11, 13–15). The GBP coatomer has thus emerged as a central hub for intracellular host defense and innate immune signaling in humans.

To visualize real-time coatomer formation by OMX-SR we deleted endogenous GBP1 in human HeLa CCL2 cells via CRISPR-Cas9 and replaced it with a functional mRFP-GBP1 reporter (GBP1^-^/^-RFP-GBP1^) expressed at physiological levels. When infected with fluorescent *Stm^EGFP^*, mRFP-GBP1 completely encapsulated individual bacilli over a ∼60-180 second time-period (**Fig. 1A**; **Movie 1**). Comprehensive coating was also observed in single-molecule W-4PiSMSN imaging of endogenous GBP1 and GBP2 on the bacterial surface in IFN-*γ*-activated GBP1^+^/^+^ cells (**Fig. 1B**; **Movie 2**). Sub-diffraction and kinetic techniques enabled average coatomer size estimates of 29,542 ± 5,156 GBP1 molecules per bacillus assembled at a rate of ∼103 ±11.6 molecules.sec^-1^ (**fig. S1A,B**).

**Fig. 1.**
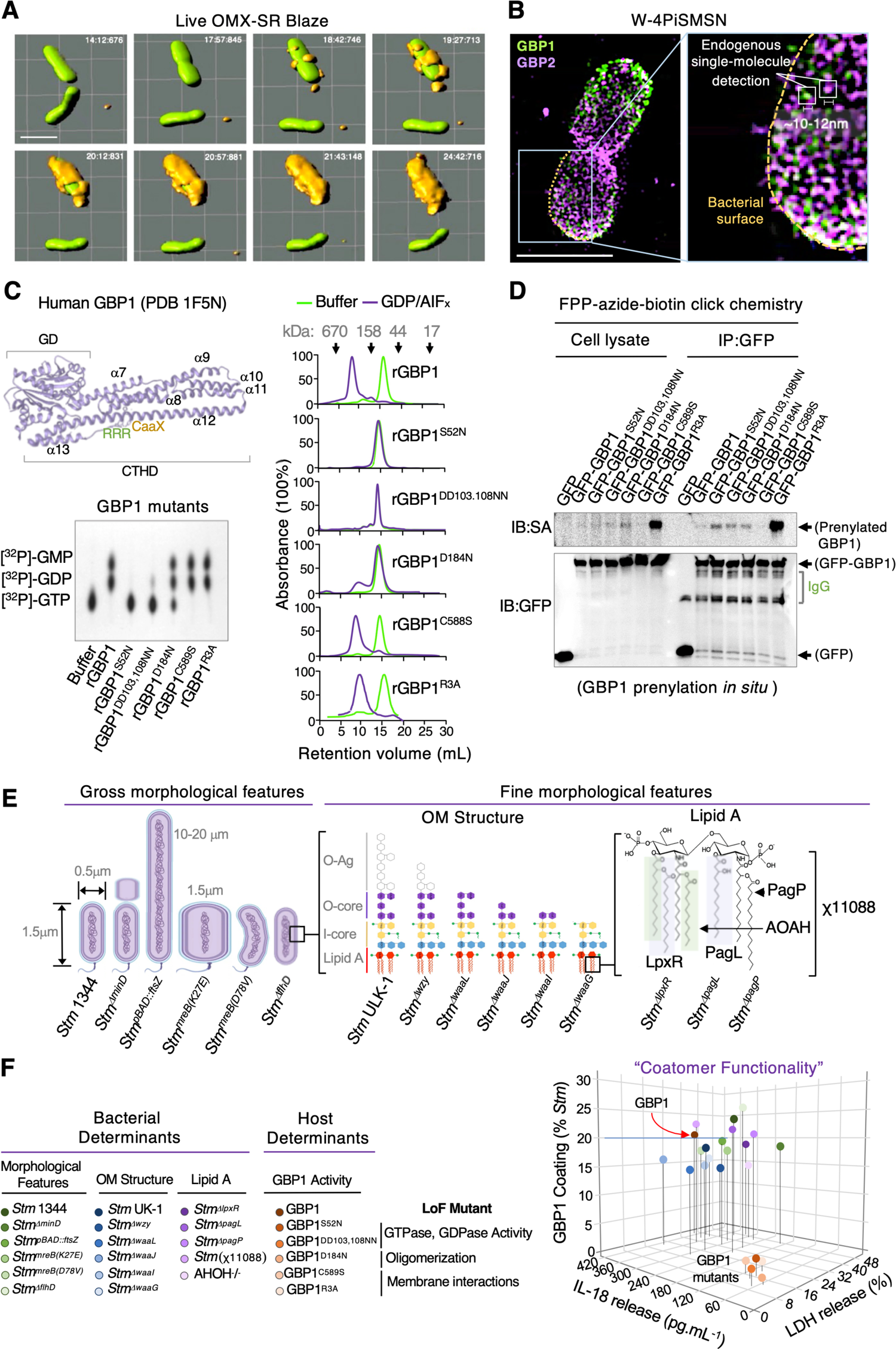
Human GBP1 coatomer kinetics and functional determinants *in situ*. (**A**) Live OMX-SR Blaze imaging that spans full encapsulation of EGFP-expressing *Stm* by RFP-GBP1 in IFN-*γ*-activated HeLa cells. Volume rendering via Imaris software. Scale bar, 2μm. (**B**) W-4PiSMSN nanoscopy of cytosolic *Stm* coated by endogenous GBP1 and GBP2 detected at single molecule resolution 2 hours post-infection. Scale bar, 1μm. (**C**) (Top) Bidomain structure of human GBP1 (PBD 1F5N, crystalized in the presence of GMPPNP) depicting catalytic G-domain (GD) and CTHD. Individual CTHD *α*-helices are denoted along with polybasic patch and farnesylation motif. (Bottom) Thin-layer chromatography of ^32^[P]-GTP hydrolysis products for recombinant GBP1 and its mutants. (**D**) Prenylation profile of EGFP-GBP1 and its mutants in HEK-293E cells detected using farnesylpyrophosphate (FPP)-azide-biotin click chemistry coupled to anti-GFP immunoprecipitation. SA, streptavidin-HRP. IgG, immunoglobulin heavy chains. (**E**) *Salmonella* and GBP1 mutants used to examine determinants of coatomer function. χ11088, *Stm^ΔlpxRΔPagLΔPagP^* triple mutant. (**F**) Coatomer “functionality” depicted by *Stm* coating and downstream IL-18 release or cell death in IFN-*γ*-activated wild-type or mutant HeLa cells infected with different bacterial strains (see color-coded key). Mean values shown. Standard deviations omitted for clarity. Representative of 3-5 independent experiments.

Such rapid kinetics required massive GBP1 co-operativity involving sequential hydrolysis of GTP and GDP for nucleotide-dependent self-assembly on the bacterial surface as revealed via GBP1 loss-of-function mutants. N-terminal GTPase (GBP1^S52N^), GDPase (GBP1^DD103,108NN^) and oligomerization (GBP1^D184N^) mutants interfered with coatomer formation and downstream IL-18 release plus pyroptotic cell death (as LDH release) in stably reconstituted GBP1^-^/^-^ cells (11) (**Fig. 1C to E**). These mutants were still post-translationally prenylated at a C-terminal CaaX motif (**Fig. 1C**) which may otherwise help anchor GBP1 to the bacterial outer membrane (OM). Indeed, mutating the CaaX box (GBP1^C589S^) blocked both C-15 farnesylation and coatomer attachment inside human cells (**Fig. 1D,E**). It did not, however, interfere with nucleotide-dependent self-assembly as shown using a transition state analog, GDP plus aluminium fluoride (AIFx), in recombinant protein assays (**Fig. 1C**). Thus, GBP1 mutants uncoupled OM attachment from subsequent polymerization, revealing distinct steps in coatomer formation during immunity to Gram-negative infection.

OM anchorage also required a polybasic patch (amino acids 584-586) resembling lipid-binding motifs in small H-Ras GTPases (19) within the GBP1 C-terminal *α*-13 helix (21; **fig. S1C**). Alanine-scanning mutagenesis of all three arginines (GBP1^R3A^) ablated coatomer formation and impacted downstream cytokine plus cell death signaling (**Fig. 1C-D**; **fig. S1D**). Because GBP1^R3A^ was heavily farnesylated the loss of coatomer assembly was not due to R3A substitution interfering with lipidation of the nearby CaaX motif (**Fig. 1D**). Instead, it appears GBP1 farnesylation is necessary but not sufficient for OM anchorage, requiring a second site to stably engage the OM and help retain it on the bacterial surface.

The bivalent nature of this C-terminal anchor was reinforced in lipopolysaccharide (LPS)-binding profiles for these two GBP1 mutants purified to homogeneity from GBP-deficient human embryonic kidney (HEK) cells to ensure correct farnesyl linkage and post-prenyl processing (21).

LPS is the major constituent of Gram-negative OMs (∼75% in *Stm*; 22) and recently reported to bind human GBP1 (14, 15). We found *Stm* LPS captured farnesylated FLAG-tagged GBP1 in fluorescence anisotropy assays at physiological pH (*K_d_*, ∼3.971μM), whereas it failed to capture non-farnesylated FLAG-GBP1^C589S^, unassembled catalytic mutants, or farnesylated FLAG-GBP1^R3A^ (**fig.S1E**). GBP1 multimerization together with the farnesyl moiety therefore appears critical for *Stm* LPS engagement with the arginine patch strengthening these interactions electrostatically to maintain a stable coat (14–15). This stability was in some cases extraordinarily long-lived: live imaging revealed a single GBP1 coat can persist for up to 2-3 hours within the human cytosol (**fig.S1E**). Thus, thousands of GBP1 molecules generate a highly durable signaling platform once anchored through initial farnesyl and polybasic contacts to the bacterial OM.

### Host and bacterial determinants of GBP1 coatomer assembly

Coatomer construction required GBP1 C-terminal attachment and catalytic activities to polymerize over the entire *Salmonella* surface. Whether microbial activities also influence this process was examined by engineering 14 *Stm* strains differing in size, shape, motility, and OM composition. *Stm* isolates that were longer (*Stm^pBAD-ftsZ^*, up to 20μm), wider (*Stm^MreB K27E^* mutants, ∼2 μm diameter) or smaller (*Stm^ΔminD^* minicells 250-300nm diameter arising from aberrant septation) still recruited endogenous GBP1 in IFN-*γ*-primed cells to activate IL-18 release and pyroptotic cell death (**Fig. 1E,F**). Bent *Stm^MreB D78V^* mutants likewise mobilized this pathway (23, 24)(**Fig. 1E,F**). Hence microbial cell size, division or curvature did not seem to influence GBP1 coatomer formation to generate an innate immune signaling platform. Flagellin-expressing (25) and flagellin-deficient *Stm* (*Stm^ΔflhD^*) were both targeted by GBP1, ruling out motility or bacterial immobilization as a cue to begin coat formation.

We therefore turned to the OM itself. Gram-negative bacteria harbor long polysaccharide LPS chains forming a divalent cation-crosslinked barrier that is impermeable to hydrophobic solutes (26). The LPS moiety consists of O-antigen polysaccharide, outer core galactose- and inner core heptose- and Kdo-enriched saccharides, and a lipid A module with multiple acyl chains embedded at the base by electrostatic and hydrophobic interactions (**Fig. 1E**). Isogenic *Stm* mutants with progressively shorter LPS chains (generated by inactivating enzymes at successive steps of the LPS biosynthetic pathway; 12, 27)(**fig. S4A**) revealed OM truncations in *Stm^Δwzy^*, *Stm^ΔwaaL^*, *Stm^ΔwaaJ^*, *Stm^ΔwaaI^* or *Stm^ΔwaaG^* did not block GBP1 coatomer formation and downstream innate immune signaling (14) (**Fig. 1E,F**).

Beneath these truncations we modified the final lipid A module, which is positioned at the base of the OM where it interacts with the underlying phospholipid inner leaflet; lipid A is recognized and directly bound by caspase-4 (28). Mutations in *Stm* LpxR or PagL (that remove a 3′-acyloxyacyl moiety or single *R*-3-hydroxymyristate chain, respectively), or PagP (that palmitoylates the hydroxymyristate chain; 29) failed to prevent GBP1 coating and caspase-4 activation (**Fig. 1E, F**). CRISPR-Cas9 deletion of human acyloxyacyl hydrolase (AOAH^-^/^-^), which is expressed at low levels in HeLa CCL2 cells and removes secondary acyl chains from lipid A as a deactivation mechanism (30), also had no effect on coatomer-dependent signaling (**Fig. 1E, F**). Thus, functional coat formation on *Salmonella* was primarily governed by host GBP1 activities rather than lipid A modifications or other microbial determinants *in situ*. Human GBP1 still targeted *Stm* irrespective of size, shape, motility, or OM composition; the latter spanned LPS chains of different length, charge, and chemical structure. Such broad ligand promiscuity may help GBP1 combat Gram-negative pathogens that modify their LPS moiety in an attempt to evade innate immune recognition and antimicrobial killing (31).

### Human GBP1 coatomer assembles a 6-member platform for cytosolic LPS recognition

Broad multivalent GBP1 interactions with the bacterial surface provide a stable platform to recruit downstream partners for innate immune signaling (11-12,15). Our previous work found GBPs2-4 and caspase-4 as part of this GBP1 signaling platform in primary human intestinal organoids and cervical epithelial cell lines (11-12,20). Here stable CRISPR-Cas9 deletions corroborated their importance with endogenous caspase-4 auto-proteolysis (denoted by the active p30 subunit), IL-18 release and pyroptotic cell death (as LDH release) significantly diminished in IFN-*γ*-activated GBP1^-^/^-^, GBP2^-^/^-^, GBP3^-^/^-^ and GBP4^-^/^-^ single knockout as well GBP1^-^/^-^2^-^/^-^ double knockout cells (**fig. S2A**). *En bloc* removal of the entire 335kb human *GBP1-7* cluster via genome engineering on chromosome 1q22.2 (GBP*^Δ^*^1q22.2^) yielded almost complete loss of downstream signaling, phenocopying CASP4^-^/^-^ and GSDMD^-^/^-^ cells (**fig. S2B**).

*GBP* gene cluster deletion in the septuple GBP*^Δ^*^1q22.2^ mutant provided a unique tool to reconstitute the entire coatomer on an empty background and test if GBP1 is the critical organizer of this hierarchical complex on the same bacilli *in situ*. Here we included full-length GSDMD as a natural caspase-4 substrate with potential bactericidal activities (32) to see if it is brought into this new supramolecular platform as well. Each coatomer component was fused to 1 of 7 fluorescent proteins (mAzurite, mSapphire, pmTurqouise2, pmEmerald, pmVenus, pmOrange, pmCardinal, pmIFP24; E_x_/E_m_ range, 384/450-684/708nm) to identify compatible combinations for reconstituting GBP*^Δ^*^1q22.2^ cells, since exchange-PAINT lacked appropriate antibodies for this spectral array and detection of endogenous proteins (**fig. S2B,C**). Our color-coded orthogonal matrix (COATOMER_450-708_) successfully resolved 5-color objects to yield a complete signaling platform with GBPs1-4 plus caspase-4 or GSDMD all coating the same bacilli after 90-120 min of infection (**fig. S2C,D**).

Remarkably, this multicolored coat was completely lost if GBP1 was omitted, indicating GBP1 establishes the entire signaling cascade at the outset (11, 15) (**fig. S3A,B**). Indeed, the COATOMER_450-708_ assay found human GBP1 was obligate for GBP2, GBP3, GBP4 and caspase-4/GSDMD simultaneously sharing the same bacterial surface. Excluding caspase-4 had no effect on GBPs1-4, placing them upstream, but largely blocked full-length GSDMD targeting, positioning it downstream in this 6-member signaling cascade as part of two-step hierarchical model (**fig. S3B,C**). Notably, N- or C-terminal GSDMD fragments mimicking the processed substrate failed to be recruited (**fig. S3C**). Hence caspase-4 brings only full-length GSDMD to this location where it is cleaved once the protease becomes activated by lipid A; this was also evident in CASP4^-^/^-^ cells where active caspase-4 restored GSDMD targeting (**fig. S3D**). Tracking the other major caspase-4 substrate in this pathway, pro-IL-18, found it failed to be recruited onto the coat (data not shown). Hence, GSDMD is a new component of the GBP1-4/CASP4 signaling complex. Its interaction with caspase-4 can likely occurs before reaching the GBP platform as shown using GBP*^Δ^*^1q22.2^ cells (**fig. S3E**).

This two-step hierarchical model was further supported by LPS binding profiles (**fig. S4A**). Post-translationally modified FLAG-GBP1, -GBP2, -GBP3, -GBP4 or -Caspase-4^C258A^ (to avoid auto-proteolysis; 28) purified from human cells and free of bacterial contaminants found GBP1 (*K_d_*, ∼3.776μM) and Caspase-4^C258A^ (*K_d_*, ∼313nM) strongly interacted with *Stm* LPS as major OM binding proteins (14–15, 28) (**sfig. 4B**). GBP1 therefore serves as the principal and initial organizer of this multiprotein complex atop the coated bacterium *in situ* (11, 15). Its assembly facilitates LPS lipid A recognition by caspase-4 that can recruit GSDMD as a subsequent step to activate cell death and cytokine release downstream.

### Bacterial sensing of the GBP1 coat alleviates periplasmic LPS accumulation *in situ*

How does the GBP1 coatomer promote LPS recognition by caspase-4, especially since lipid A is buried at the base of the OM? We probed LPS release and OM stress responses in live bacteria *in situ* (**Fig. 2A-F**). First, copper (Cu^2+^)-free CLICK chemistry was used to label *Stm* LPS with Alexa Fluor attached via Kdo-azide derivatives adjacent to lipid A within the inner core (**Fig. 2A**). Here anti-*Salmonella* O-antigen antibody was used in conjunction to verify the KDO-Alexa Fluor signal which decreases during its transit to the cytosol. We likewise incorporated fluorescent D-alanine into the L-Ala-D-*meso*-diaminopimelate-D-Ala-*O*-Ala pentapeptide via labelling of the underlying peptidoglycan scaffold in metabolically active bacteria (33)(**Fig. 2A**). 3D structured illumination microscopy (SIM) found complete GBP1 coating triggered release of LPS but did not appear to disturb the *Stm* peptidoglycan layer within the cytosol of IFN-*γ*-activated HeLa cells (**Fig. 2B, F**). LPS liberation was significantly diminished in GBP1^-^/^-^ epithelia that also exhibited defects in caspase-4 activation (**Fig. 2F**), confirming GBP1 can promote caspase-4 ligand availability *in situ*. Other exteriorized structures such as flagella were still evident on GBP1-coated bacteria (**Fig. 2B**); hence GBP1 primarily affected LPS release, leaving the rigid PG scaffold and flagellar apparatus intact.

**Fig. 2.**
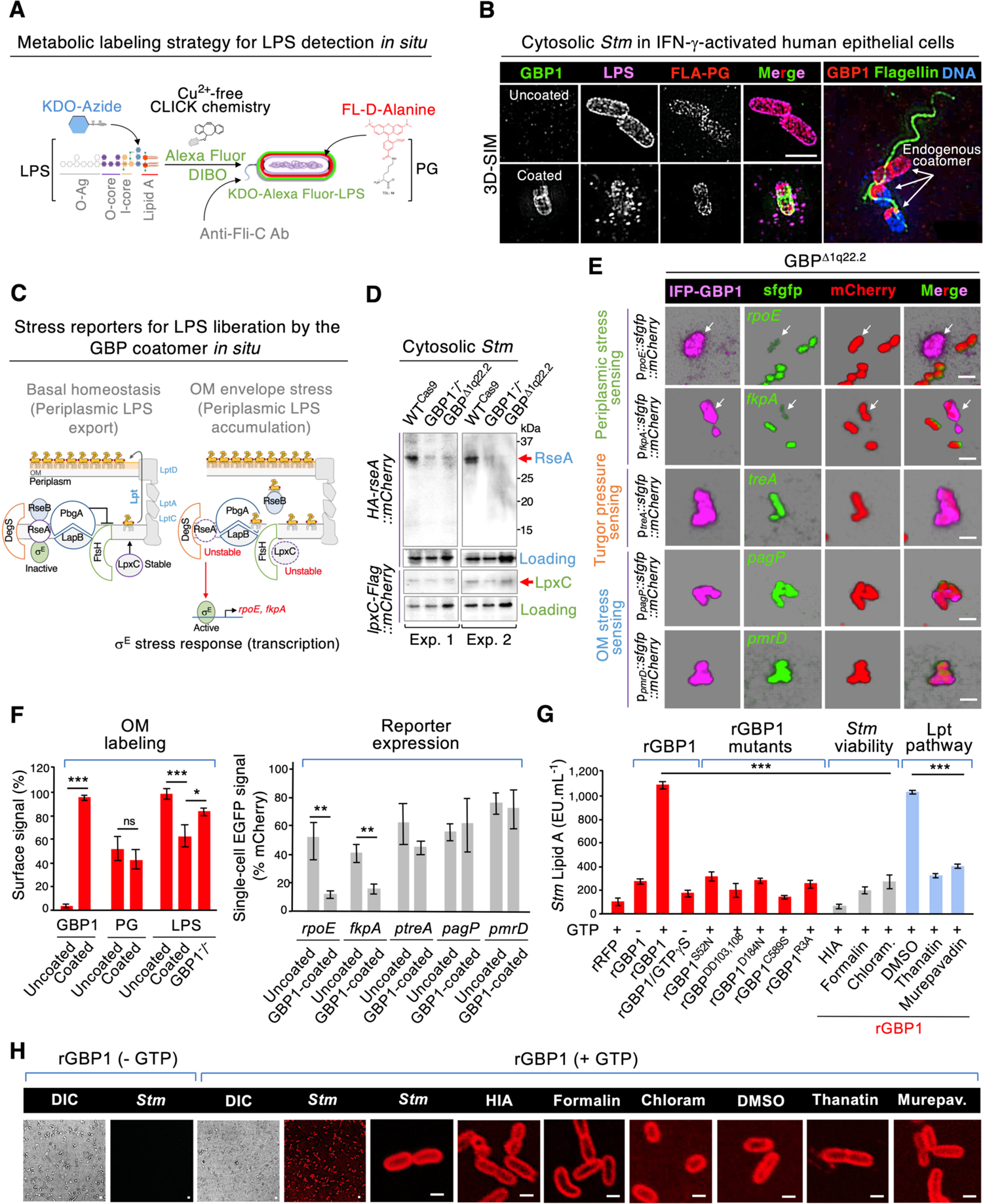
The GBP1 coatomer is sensed by *Salmonella* for LPS release *in situ* and in cell-free systems. (**A**) Fluorescent labeling strategy for LPS using copper-free click chemistry with a dibenzocyclooctynol (DIBO) alkyne intermediate. Incorporation of HCC-amino-D-alanine (HADA) was used for peptidoglycan. KDO, 3-deoxy-D-manno-octulosonic acid. (**B**) 3D structured illumination microscopy (SIM) of endogenous GBP1 coating detected by anti-GBP1 antibody in HeLa cells activated by 1,000U IFN-*γ*for 18h. Pseudo-colored images collected at 2 hours post-infection with pre-labelled *Stm* at MOI of 20. Endogenous flagellin detected with anti-Fli-C antibody and LPS with anti-Sal-O antibody for verification. Scale bar, 2μm. (**C**) Depiction of periplasmic LPS stress reporters that undergo degradation for activation the *σ*^E^ stress response. (**D**) Immunoblot of cytosolic bacterial lysates expressing hemagglutinin (HA)-fused RseA or Flag-LpxC retrieved at 2-hour post-infection from IFN-*γ*-activated wild-type, GBP1^-^/^-^ or GBP*^Δ^*^1q22.2^ cells shown in two independent experiments. Loading controls included mCherry expressed by a constitutive *rpsM* promoter as part of a dual reporter system (for RseA) and a non-specific band (nsb) detected by the same anti-Flag antibody as a more sensitive control for low *in situ* LpxC-Flag expression. (**E**) Triple-color scanning confocal microscopy of *Stm* strains expressing different stress reporters inside GBP*^Δ^*^1q22.2^ cells complemented IFP-GBP1 to test for sufficiency. Images collected at 3 hours post-infection. Arrows, reduced reporter expression in GBP1-targeted bacteria. (**F**) Quantitation of metabolic labeling and reporter expression. (Left) Percentage of pre-labelled *Stm* with intact signal measured by InCell software. PG labeled ∼50% of the total surface *in situ* as unlabeled PG becomes incorporated into dividing bacteria. *n* = 95 bacilli examined from 3 independent experiments. (Right) Single-cell ratiometric EGFP:mCherry reporter expression at 3 hours p.i. (fluorescence intensity [MFI] GFP signal as percentage of mCherry signal within the same individual bacillus). *n* = 45-60 bacilli per group from 4 independent experiments. (**G**) Soluble lipid A release detected by LAL in coatomer reconstitution assays (triplicate ± SD) in the presence of recombinant GBP1 with or without GTP substrate or a non-hydrolyzable GTP*γ*S analog. rGBP1 mutants did not coat live bacilli. HIA, heat-inactivated; chloram., chloramphenicol; DMSO, dimethyl sulfoxide (control for DMSO-solubilized inhibitors). Representative of 3-6 independent experiments. (**H**) Confocal imaging of cell-free *Stm* 1344 coated for 60 min with rRFP-GBP1 ± GTP. Treatments shown above. Scale bar, 1μm. * p<0.5, ** p<0.01, ***p<0.001. Significance values using Students t-test (F) or one-way ANOVA with Holm-Sidak post-hoc test (G).

The OM of Gram-negative bacteria is an essential load-bearing element that deforms with stretching, bending and indentation forces (34). LPS liberation could thus arise from constriction applied by GBP1 acting as a dynamin-like mechanoenzyme. Measuring bacterial turgor pressure via plasmolysis, however, proved insensitive inside human cells. As an alternative strategy, we used bacterial reporters of turgor pressure and LPS homeostasis to inform us how bacteria sense the presence of the GBP1 coat inside the human cytosol (**Fig. 2C-E**). Loss of turgor in *Stm* induces expression of TreA, a periplasmic trehalase that hydrolyses the osmoprotectant trehalose into two glucose molecules for osmotic repair (35). Loss of LPS homeostasis can yield dual degradative signals. First, a regulatory protein RseA is degraded by an inner membrane protease, DegS, to help release the cytoplasmic σ^E^ transcription factor subunit for stress-response gene expression (36). Among these stress-response genes are isomerases and chaperones that refold proteins in the OM and LPS transport pathways (Lpt) (26). Second, degradation of the LPS assembly enzyme, LpxC, by the protease FtsH, stops *de novo* synthesis to prevent toxic build-up of LPS intermediates within the periplasm (37)(**Fig. 2C**).

We engineered *Stm* strains with arabinose-responsive HA-tagged RseA or FLAG-tagged LpxC to ensure bacteria expressed these reporters in the host cytosol to assess their fate when coated by GBP1. Unexpectedly, GBP1 coating protected RseA from degradation in IFN-*γ*-primed HeLa cells but was heavily degraded in GBP1^-^/^-^ or GBP*^Δ^*^1q22.2^ cells that lack the ability to produce a GBP coat; LpxC proteolysis was less sensitive as a reporter (**Fig. 2D**). This result was confirmed by single *Salmonella* EGFP expression from *rpoE* or *fkpA* promoters as σ^E^ targets downstream of RseA degradation (**Fig. 2C**). Uncoated *Stm* had robust *rpoE-* or *fkpA*-driven EGFP expression, whereas up to 75% of GBP1-coated *Stm* did not, despite bacterial co-expression of RFP from a constitutive ribosomal *rpsM* promoter as an internal control (**Fig. 2E,F**). In contrast, EGFP expression from the *treA* promoter or PhoP-PhoQ-driven *pmrD* and *pagP* promoters were equally robust in coated and uncoated bacteria (**Fig. 2E,F**). Hence GBP1 encapsulation appeared to reduce σ^E^ stress responses *in situ*. Insulating bacteria from certain types of OM stress prevents toxic accumulation of off-target intermediates within the periplasm (36) and maintains an uninterrupted supply of newly synthesized LPS to the bacterial surface via the Lpt pathway (26). Here unanchored lipid A may be bound by caspase-4 following its recruitment by GBP1 (11).

### A reconstituted coatomer confirms GBP1-dependent periplasmic LPS release

We next measured unanchored (soluble) lipid A release in a cell-free coatomer assay to test if GBP1 was indeed sufficient to promote LPS liberation. This simplified setting eliminated other host cell sources of environmental stress and allowed soluble lipid A detection that cannot be measured *in situ* due to whole bacterial cell contamination of the host cytosol. Incubation of farnesylated recombinant RFP-GBP1 (**fig. S5A,B**) with axenic *Stm* led to >95% of bacteria becoming fully coated within 60 minutes after addition of GTP substrate; encapsulation followed a highly accelerated sigmoidal curve yielding a half maximal value (“coatomer K_m_”) of 225nM and steep Hill slope of 5.22 (**Fig. 2E and fig. S5C**). Notably, this all-or-none behavior did not arise from crossing a phase transition boundary since rRFP-GBP1 does not phase separate, unlike plant GBPLs (7) (**fig. S5D**). Farnesylated RFP-GBP1^C589S^ and other non-coating GBP1 mutants identified *in situ* were absent from *Stm* even after addition of GTP, mimicking the results seen in human cells (**Fig. 2E**). Pronounced coating was evident for *Stm* irrespective of bacterial size or LPS status; comparison with Gram-negative *P. aeruginosa* and Gram-positive *L. monocytogenes* also showed the latter bacteria had much lower levels of GBP1 encapsulation (**sfig.6A-C**).

Importantly, reconstituting the GBP1 coatomer triggered *Stm* LPS release as measured by limulus amebocyte lysate (LAL) assay that detects the soluble lipid A moiety. The amount of lipid A released from coated *Stm* exceeded the *K_D_* range of caspase-4 (28) yet comprised < 1% of the total LPS present (based on 2 x 10^6^ molecules of LPS per *Stm*; 22). Hence the GBP1 coatomer does not seem to instigate wholesale disruption of the OM, a finding compatible with the diminished stress profiles of GBP1-coated bacteria *in situ* and recent reports showing GBP1 on its own does not kill intact Gram-negative bacteria in human cells or cell-free systems (11-12, 14-15). Instead, GBP1 insertion probably affects lateral LPS-LPS cationic interactions more subtly to compromise OM stiffness and allow passage of small antimicrobial proteins such as APOL3 (12). It also suggests LPS may arise from additional sources besides the pre-existing OM pool if milder GBP1 damage is common.

We tested this idea in live metabolically active bacteria versus those rendered non-viable by heat-inactivation, formalin-fixation, or chloramphenicol treatment. LPS release was essentially lost in the latter group despite being equally well coated by rRFP-GBP1 (**Fig. 2G,H**). Thus, only viable *Stm* released significant amounts of lipid A in response to GBP1 encapsulation in these cell-free assays. This result hints at bacteria sensing the presence of the GBP1 coat to maintain the LPS export pathway as seen with bacterial reporters inside human cells. This likelihood was further reinforced by inhibitors of the Lpt pathway (26). Live bacteria pre-treated with thanatin or murepavadin to block the ascending LptC and LptD subunits of the Lpt barrel complex, respectively, interfered with lipid A release from GBP1-coated bacteria (**Fig. 2C,G** and H). Thus, anchorage of human GBP1 to the bacterial surface not only stabilizes the caspase-4-GSDMD assembly platform but seems to maintain active LPS export from the underlying periplasmic space. Importantly, GBP1 alone was sufficient for LPS release, irrespective of whether LPS originated from periplasmic stores or was directly dislodged from the outer leaflet by its insertion into the OM.

### A minicell system enables ultrastructural studies of the native GBP1 coatomer

Our reconstituted coatomer system allowed us to delineate how GBP1 inserts into the outer leaflet at a molecular level for triggering OM and periplasmic LPS responses. Here GBP1 attachment could be viewed with powerful quasi-atomic tools such as cryo-ET. We engineered bacterial minicells for coating with rRFP-GBP1 since they are considerably smaller than isogenic rod-shaped bacilli. The reduced sample thickness of minicells improves resolution in cryo-ET samples (23).

Minicells arise from abnormal asymmetric cell division (**Fig. 3A**); we used arabinose-induced expression of the septation gene, *FtsZ*, to generate *Stm* minicells of the appropriate size (∼150-300nm) and which were sufficiently robust for isolation by differential centrifugation along with outer membrane vesicles (OMVs) (**Fig. 3B**). *FtsZ* over-expression was introduced onto a *waaG*-deficient background lacking the LPS O-antigen and outer core segment (*Stm^ΔwaaG::pBAD-ftsZ^*) because recent work found these segments possess unstructured density that interfere with sub-tomogram averaging (38). O-antigen and outer core segments are also dispensable for GBP1 coatomer attachment and downstream signaling inside human cells (**Fig. 1G**). Our dual genetic strategy was therefore devised to limit sample noise during the acquisition of tilt images by cryo-ET.

**Fig. 3.**
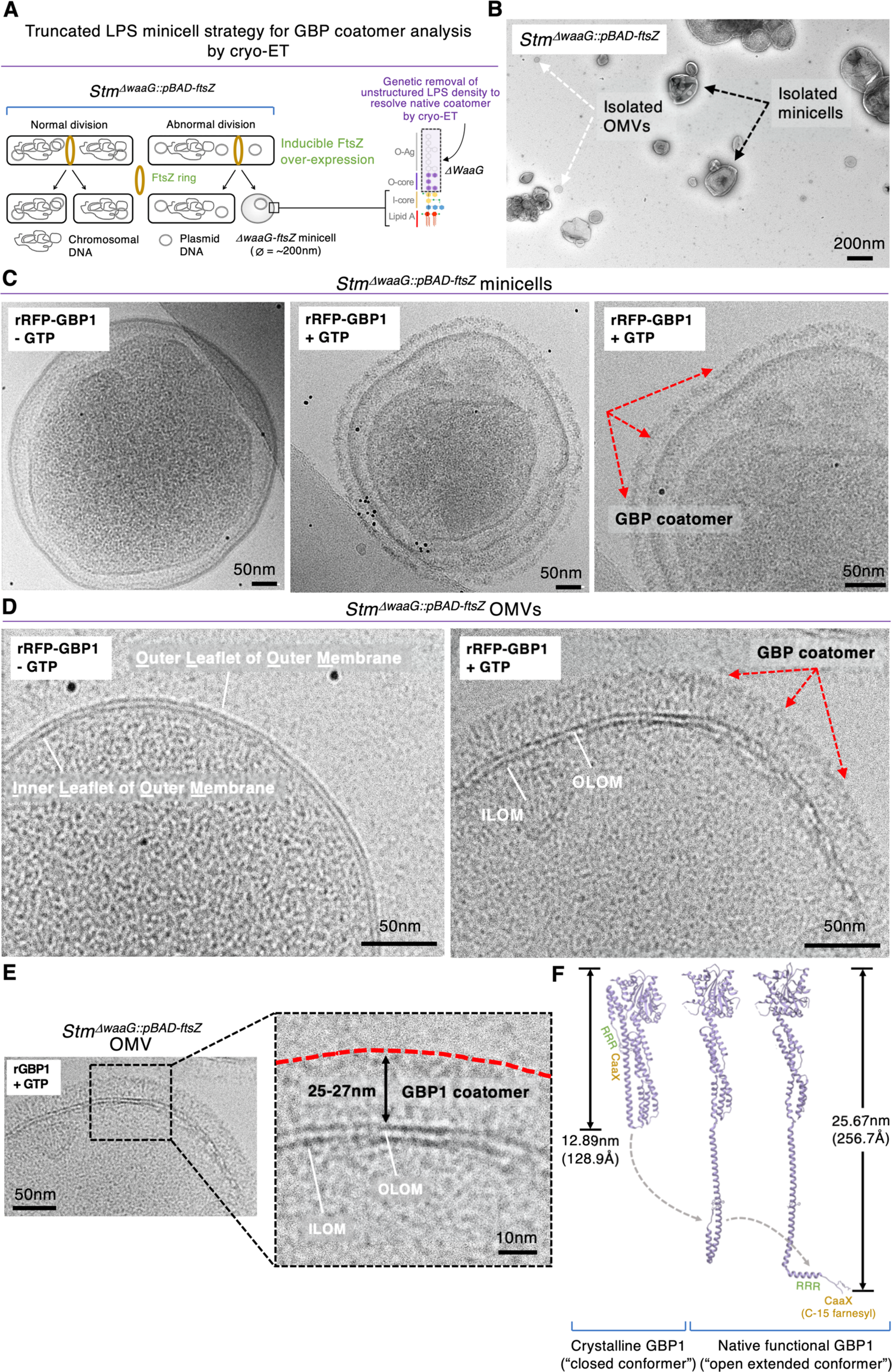
*In vitro* reconstitution of GBP1 coating assembly on the *Salmonella* minicell observed by cryo-EM. **(A)** Strategy for constructing *Stm^ΔwaaG::pBAD-ftsZ^* minicells missing LPS O-antigen and outer core for improved cryo-ET resolution. **(B)** Representative negative stain images showing isolated fractions have both outer membrane vesicles (OMVs) and minicells. **(C)** GTP-dependency of hGBP1 coatomer formation on the *Stm^ΔwaaG::pBAD-ftsZ^* minicell surface shown in 200kV cryo-EM images. Black dots, 6nm diameter fiducial beads. **(D)** GTP-dependency of hGBP1 coatomer formation on the *Stm^ΔwaaG::pBAD-ftsZ^* OMV surface shown in cryo-EM images. **(E)** Estimated ∼25-27nm GBP1 coatomer length *Stm^ΔwaaG::pBAD-ftsZ^* OMVs. OLOM, outer leaflet of outer membrane. ILOM, inner leaflet of outer membrane. **(F)** Dynamic GBP1 modeling based on initial size estimations from cryo-EM.

Cryo-EM at 200kV revealed successful reconstitution of the GBP1 coatomer complex on *Stm^ΔwaaG::pBAD-ftsZ^* minicells and OMVs in comparative samples with rRFP-GBP1 ± GTP (**Fig. 3C,D**). Initial EM measurements found GBP1 spans ∼25-27nm tightly juxtaposed over the entire bacterial surface (**Fig. 3E**). Crystallization of human GBP1 bound to the GTP analogue, GMPPNP (PDB 1F5N), is half this length (12.89nm, 128.9Å; 39), with the final *α*13 helix collapsed back onto the *α*12 C-terminal segment (**Fig. 3F**). Hence farnesyl anchorage to the OM would require either head-to-head or head-to-tail dimers spanning 257.8 Å to fit the EM measurements, or 4-6 GBP1 monomers horizontally arrayed atop one another and perpendicular to the outer leaflet (39). Alternatively, GBP1 may undergo dynamic C-terminal extension to present the fully unhinged C-15 farnesyl group to the OM as modeled in our *ab initio* computer simulations (**Fig. 3F**). This predicted conformer opens to 256.7Å. Each potential configuration was probed by cryo-ET to discern how GBP1 directly operates on the bacterial surface under native conditions.

### Thousands of extended 260Å GBP1 conformers insert into the bacterial OM

To view the native GBP1 coatomer on *Stm^ΔwaaG::pBAD-ftsZ^*, we used 300kV field-emission instrumentation to collect high contrast images in dose-fractionated mode. In some images rRFP-GBP1-coated OMVs were also detected (**Fig. 4A**). Iterative classification of 20,148 coatomer particles yielded 6 initial classes where outer and inner leaflets of the OM membrane was evident (**Fig. 4B**). Further refinement of individual protomers necessitated user-side scripting based on the Pyseg package to accommodate both flexibility of the GBP1 structure and its small 67.8kDa size. This strategy enabled us to align stack files for 3D segmentation of the human GBP1 coatomer complex at higher pixel/resolution ratios on the *Salmonella* OM (**Fig. 4C**). Solving this massive complex to 31.07Å demarcated a globular G-domain and fully extended C-terminal helical domain (*α*7-*α*13 helices) that could be pseudo-modelled, confirming both the orientation and dynamic opening of GBP1 to insert near the lipid A module (**Fig. 4D,E**). These open GBP1 conformers were ∼260Å long and rough extrapolation gave between 12,000-15,000 molecules per minicell, yielding a theoretical ∼800MDa-1.02GDa structure. Thus, cryo-ET provided an initial quasi-atomic view of the native coatomer on the bacterial surface where thousands of GBP1 molecules stretch their C-terminal helical domain to ∼260Å for anchoring the farnesyl tail and insertion of the nearby polybasic patch. Such flexibility for GBP1, particularly *α*13, has been alluded to in sub-millisecond atomistic H-REMD simulations (40). However, the native GBP1 conformer has never been resolved empirically before, especially on a physiological substrate like the bacterial OM.

**Fig. 4.**
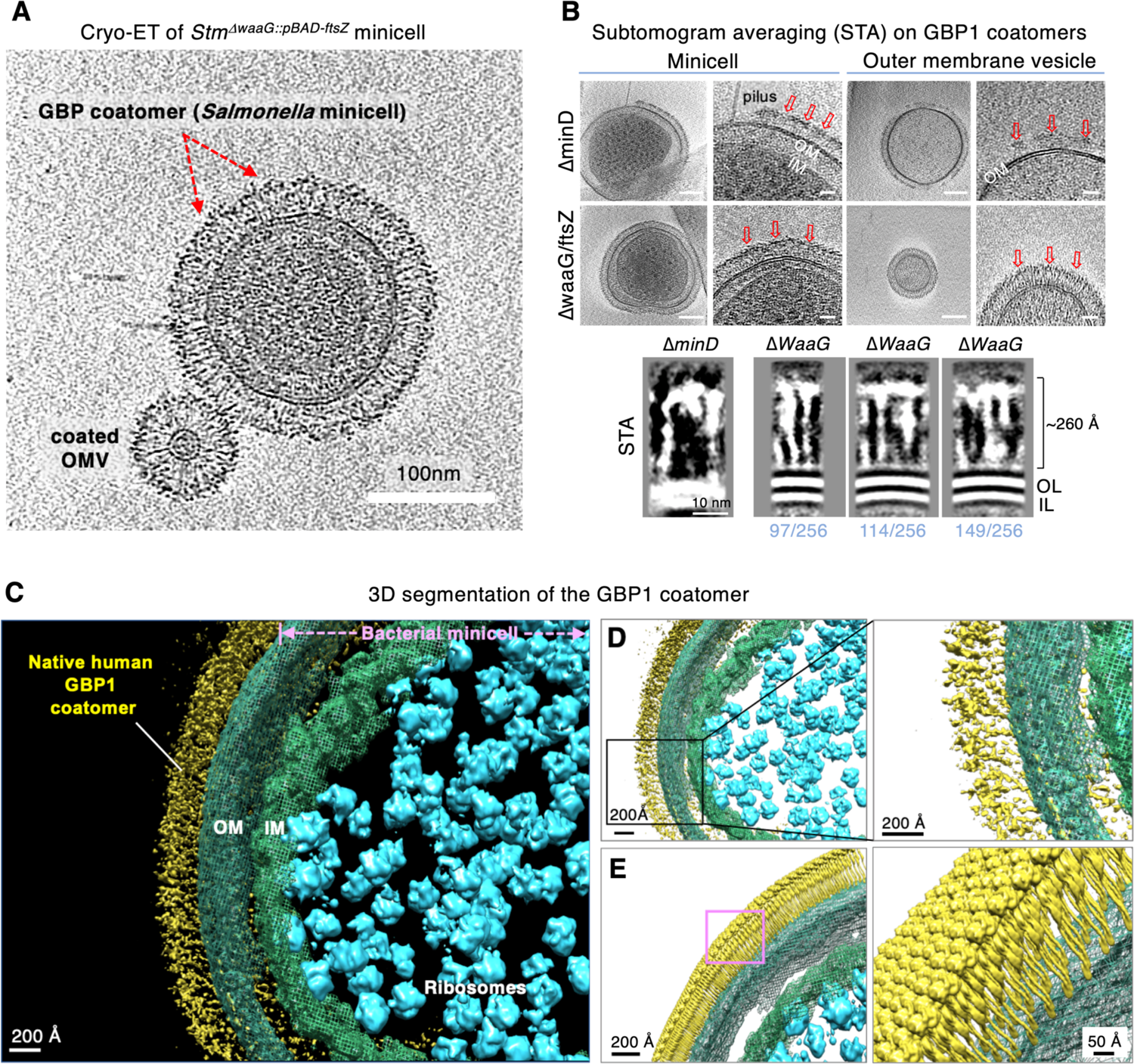
Extended GBP1 conformer observed in a functional state by cryo-ET and sub-tomogram averaging. **(A)** Representative tomographic slice of *Stm^ΔwaaG::pBAD-ftsZ^* minicell completely coated with rRFP-hGBP1. Extended GBP1 conformers are visible in coated OMVs. **(B)** (Left) Tomographic slice images of minicells or OMVs isolated from *Stm^ΔminD^* and *Stm^ΔwaaG::pBAD-ftsZ^* for comparison. (Right) Corresponding zoom-in view of the reconstituted GBP1 coatomer. Red arrows depict either full or partial coatomer. OM, outer membrane; IM, inner membrane. Scale bar, 100 nm in overall image; 25nm for zoom-in view. (Bottom) Left to right, sub-tomogram average of GBP1 targeting on *Stm^ΔminD^* minicell and three different slice views of sub-tomogram average for GBP1 on *Stm^ΔwaaG::pBAD-ftsZ^* minicells. 97/256, 97th slicer of 3D volume within 256*256*256 voxels; 114/256, 114th slicer of 3D volume; 149/256, 149 slice images. STA, sub-tomogram averaging. OL, outer leaflet of outer membrane. IL, inner leaflet of outer membrane. **(C)** 3D surface rendering of the GBP1 coat using post-acquisition segmentation. OM, outer membrane; IM, inner membrane **(D)** Zoom-in view of 3D segmentation of the GBP1 coatomer reveal outermost globular density and “stalk-like” attachment underneath. Blue, bacterial ribosomes; Green, bacterial membrane; yellow, GBP1 coat. **(E)** Human GBP1 crystal structure (1F5N) was low-pass filtered to 15 Å to build a coatomer pseudo-model on the bacterial surface.

### Cryo-ET identification of the native GBP1 coatomer *in situ*

Bacterial minicells enabled resolution of individual GBP1 conformers by cryo-ET within a mesoscale complex *in vitro*. Whether these extended conformers yield a ∼25nm coatomer complex *in situ* was next examined in focus ion beam-(FIB)-milled lamella corroborated before and after by correlative light and electron microscopy (CLEM) (**Fig. 5A, B**). FIB milling typically generates lamella ∼150-300nm in thickness; this may miss coating events which, unlike >95% GBP1-encapsulated bacteria in cell-free reconstitution assays, are comparatively rare across the entire ∼2,500μm^3^ volume of an intact human HeLa cell (BNID 103725). Hence, we first tested ultrathin human U2OS osteosarcoma cells ∼200nm thick to increase coatomer events in lamellae; low *Stm* invasion rates, however, precluded their use. We therefore opted to increase bacterial size instead. *Stm^pBAD-ftsZ^* bacilli up to 20μm in length were used to infect GBP*^Δ^*^1q22.2^ cells expressing RFP-GBP1 (**Fig. 5A**). GBP*^Δ^*^1q22.2^ cells were chosen since they cannot assemble the 6-member coat (**sfig. 3A**); hence only reintroduced EGFP-GBP1 was present on the bacterial surface for cryo-ET. In addition, FACS sorting EGFP-GBP1^+^ cells for *Stm* infection (12) further enriched for these events before FIB milling (**Fig. 5A**).

**Fig. 5.**
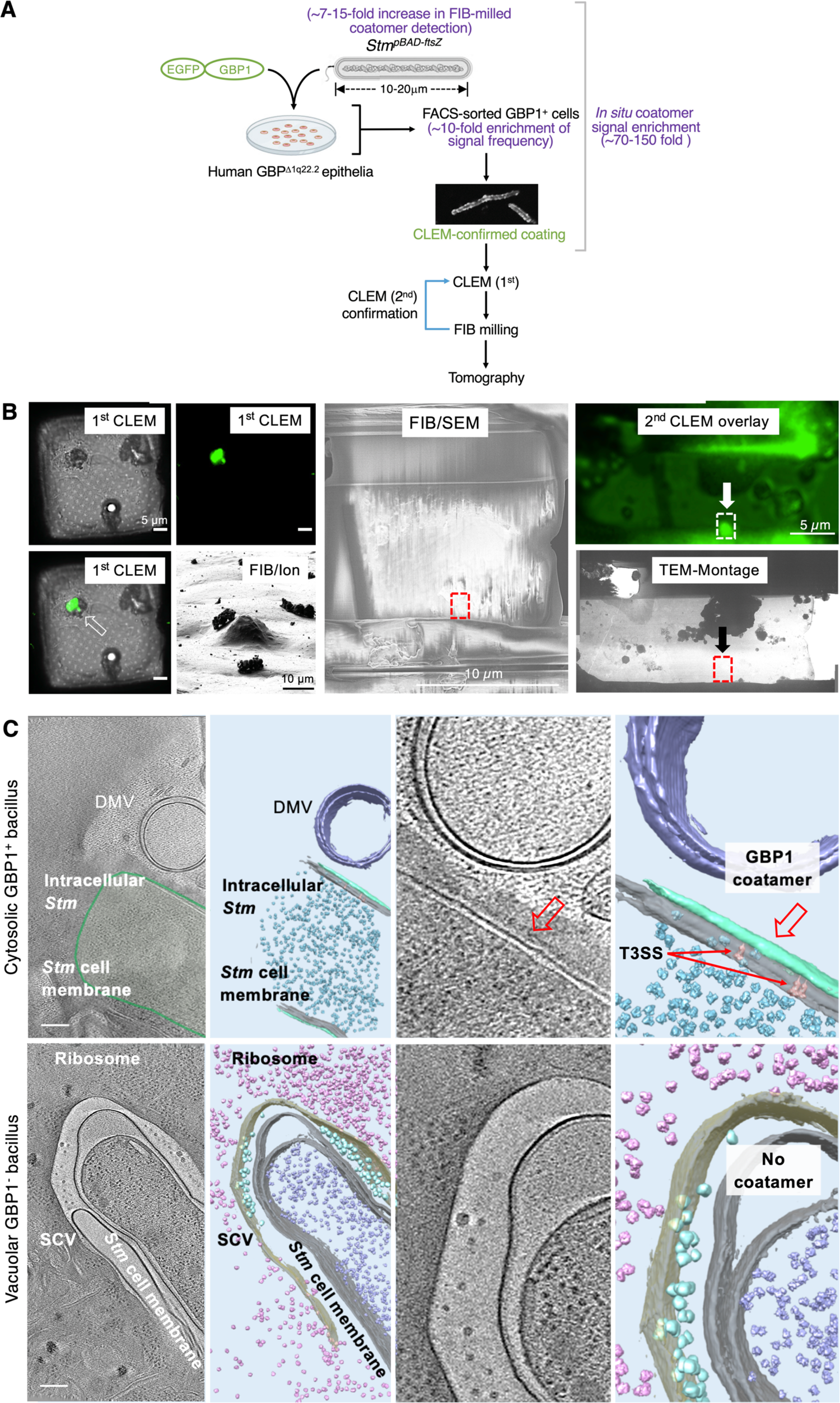
*In situ* GBP1 coatomer visualization by cryo-ET on FIB-milled and CLEM-verified lamellae. **(A)** Cryo-CLEM workflow of EGFP-GBP1 detection *in situ*. **(B)** 1^st^ Cryo-CLEM, FIB and 2^nd^ cryo-CLEM validation of EGFP-hGBP1 coating on elongated *Stm^pBAD-ftsZ^*. Arrow, milling target for EGFP-GBP1 coatomer. 1^st^ CLEM image was taken with DIC and GFP fluorescent channels. FIB/Ion, focused-ion beam image before rough milling; FIB/SEM, SEM image after fine milling; 2^nd^ CLEM overlay, cryo-CLEM observation confirming EGFP-hGBP1 fluorescent signal in the grid after fine milling of FIB; TEM-Montage; transmission EM of the milled lamellae. Dashed rectangle, area where cryo-ET tilt series were collected and matched to the top panel of tomogram in (**C**). **(C)** (Top panel) *In situ* tomographic slices with 17.2 nm thickness. Cytosolic *Stm^pBAD-ftsZ^* was targeted by EGFP-GBP1. DMV, double-membrane vesicle. Zoomed-in area at right shows GBP1 coatomer (red arrow) and bacterial type 3 secretion system (T3SS) complex. (Bottom panel) Representative tomographic slice with 17.2 nm thickness of vacuolar *Stm^pBAD-ftsZ^* lacking GBP1 coating. In these 3D segmented images, cytosolic free GBP*^Δ^*^1q22.2^ cell ribosomes are indicated in the pink; Bacterial ribosomes are segmented in the purple. SCV, *Salmonella*-containing vacuole. Scale bar, 200 nm.

This approach successfully captured EGFP-GBP1^+^ *Stm^pBAD-ftsZ^* in FIB-milled lamella (**Fig. 5B**). A delineated coat was present only on CLEM-validated cytosolic bacteria compared with *Stm* in vacuoles which are negative for GBP1 (**Fig. 5C**). *In situ* cryo-ET coatomers displayed similar length characteristics to coats in cell-free reconstitution: a solid ∼25nm GBP1 boundary around the perimeter of the OM (**Fig. 5C**). At this resolution individual 68kDa conformers could not be deciphered although our analysis reveals the coatomer can be successfully detected inside human cells after post-FIB verification by CLEM. Future studies will help refine this massive object *in situ*. It should provide mechanistic insights into how human GBP1 establishes its foothold on the LPS leaflet to trigger ligand release and polymerization over for entire Gram-negative bacterial surface once the pathogen is recognized in the cytosol.

## Discussion

Higher-order protein assemblies help amplify innate immune signaling and spatially regulate signal propagation by localizing partners at the site of ligand recognition (1, 4). Such assemblies form on the host plasma membrane, mitochondria, peroxisomes, and chloroplasts (1, 41). They also occur in the cytosol, nucleus, ER and endosomal network, in some cases yielding membrane-less condensates via liquid-liquid phase separation (42), as recently discovered for plant GBPLs during cell-autonomous defense against bacterial phytopathogens (7).

By contrast, human GBP1 builds a multiprotein complex on a completely foreign object – the Gram-negative bacterial OM. This huge nanomachine solicits GBPs2-4, caspase-4 and full-length GSDMD as part of a 6-member platform to propagate cytokine and cell death signaling in multiple human cell types (11, 15). It likewise facilitates bacterial killing by IFN-*γ*-induced APOL3 in non-immune cell populations as well (12). Early studies discovered GBPs target intracellular bacteria and physically assemble antimicrobial or inflammasome complexes (8, 9). Here the involvement of human GBPs in caspase-driven activities arose from *in silico* Hidden Markov modeling that retrieved GBP-related sequences in lower organisms sharing CARD or DED domain modularity with the human inflammasome and cell death machinery (9). This evolutionary relationship now extends to humans, mice, and zebrafish where multiple GBPs bind inflammatory caspases, NLR sensors or ASC adaptors to facilitate microbial recognition (9–11, 13).

Human GBP1 and GBP3 both co-immunoprecipitate endogenous caspase-4 yet perform distinct functions within infected cells (11). Our current study shows GBP1 can trigger periplasmic and OM lipid A release which activates caspase-4 after recruiting it to the bacterial surface. In contrast, GBP3 is dispensable for caspase-4 recruitment but may control its catalytic activity (11). Unlike GBP1, neither GBP3 nor GBP2 or GBP4 purified from human cells strongly bound *Stm* LPS; hence their functions probably differ from known LPS transfer proteins such as LBP-CD14-MD2 within the TLR4 cascade (41). Future work will clarify the precise role of these other GBPs within the assembled signalosome.

Human GBP1 is the central organizer of this supramolecular platform and is reported to coat several cytosol-invasive Gram-negative pathogens including *S. typhimurium*, *F. norvicida* and *S. flexneri^ΔIpaH9.8^* mutants to activate caspase-4 (11-15, 20, 43). Its promiscuous LPS binding properties allow it to overcome unusual lipid A alterations such a tetra-acetylation by *F. norvicida* (31). Such multivalency likely extends beyond LPS to other lipoglycan structures and probably explains why GBP1 also encapsulates some Gram-positive pathogens, mycobacteria, *T. gondii* and norovirus replication complexes in IFN-*γ*-activated human and mouse macrophages to mobilize host defense (8, 13, 20, 44). In all cases, oligomerization and farnesylation was paramount.

GBPs belong to the dynamin-like protein superfamily that exhibit high intrinsic GTPase activity (*k_cat_* ∼2-100 min^-1^), low μM substrate affinity and nucleotide-dependent self-assembly inside cells (45). We found nucleotide-driven GBP1 co-operativity gave a sigmoidal *Stm* coating curve on bacteria that steeply accelerated above 125nM, like other “prionizing” proteins where all-or-none responsivity occurs once a concentration threshold is reached (4). Here anchorage to LPS may accelerate GBP1 catalysis and oligomerization (15). Both GTPase and GDPase activities contributed to GBP1 responsivity via single GTP/GDP turnover as part of a 2-step enzymatic process (14, 46). Because hydrolysis of GTP/GDP liberates large amounts of Gibbs free energy the GBP1 coatomer conforms to a nanomachine by performing “work” in establishing this massive signaling platform.

C-terminal anchorage was likewise critical for coatomer assembly. It not only laid the physical foundation but released soluble lipid A for caspase-4 autoproteolysis. GBP1 coating unexpectedly insulated against periplasmic stress where aberrant LPS accumulation triggers the σ^E^ stress regulon (36). These stress responses occur as *Stm* runs the gauntlet of the host endolysosomal system before escaping its vacuole to enter the cytosol (12, 35). By preventing disruption of LPS homeostasis, the GBP coatomer may ensure a steady supply of lipid A via the Lpt pathway for caspase-4 activation. Moreover, turning off the downstream RpoE regulon lowers resistance to antimicrobial peptides and makes *Stm* more vulnerable to attack (47); potential bactericidal agents include IFN-*γ*-induced APOL3 known to co-operate with GBP1 (12). Thus, C-terminal attachment not only generates a stable GBP signaling platform but likely disables bacterial evasion mechanisms as well.

This C-terminal attachment relied on farnesylation and a neighboring 584-586 polybasic patch. C-15 lipidation requires sequential addition by human farnesyltransferase, tripeptide removal by CAAX carboxypeptidase (RCE1), and carboxy-group methylation by isoprenylcysteine carboxymethyltransferase (ICMT) (21). Post-prenyl processing thus brings the fully modified farnesyl tail almost adjacent (3 amino acids apart) to the triple arginine patch, creating a powerful bipartite anchor. Genetic ablation of RRR abolished *in situ* coating but not *in situ* farnesylation, whereas CaaX box mutation abolished farnesylation-based attachment but left RRR intact. Hence each was necessary and neither sufficient for bacterial encapsulation, reinforcing their combined actions for OM anchorage. Here insertion of the polybasic motif could undergo electrostatic interactions with the negatively charged PO_4_^-^ groups of lipid A and inner core saccharides (14, 15); 3D segmentation of the native GBP1 conformer positioned the C-terminal extension near both lower LPS segments in the *waaG*-*ftsZ* minicell and OMV mutant.

Cryo-ET provided us with a quasi-atomic view of native GBP1 and its supramolecular architecture directly on the Gram-negative *Salmonella* surface. Previous crystal structures of GBP1 used full-length recombinant protein produced in *Escherichia coli* to capture the monomer (apo) state ± GMPPNP, a GTP analogue, at 1.7-2.3 Å (40, 48). The N-terminal G-domain was likewise produced in bacteria and crystallized in the presence of multiple nucleotides as a dimer to 2.9 Å (46). Both full-length human GBP1 crystal structures position *α*-12 and *α*-13 helices tucked up against the *α*7-*α*11 middle domain in a folded configuration, whereas GBP1 appears fully extended with the *α*-12 and farnesylated *α*-13 helices inserting vertically into the bacterial outer leaflet under physiological conditions. Visualizing the open GBP1 conformer as part of a mature coat, especially *in situ*, reinforces the capacity of cryo-ET to give mechanistic insights into how assembled proteins behave within a natural cellular context (49).

Initial resolution of this 3D coatomer proved challenging across two scales: first, the small size of individual GBP1 monomers, and second, the massive size of the final polymer. These sizes spanned 68 to >800,000 kDa. Delineating the length and orientation of individual GBP1 molecules to 31Å among thousands of identical proteins benefitted from bacterial genetics plus recombinant protein preparation. *Stm^ΔwaaG::pBAD-ftsZ^* minicell sizes (100-200nm) helped reduce inelastic electron scattering during tomography whereas genetic removal of the LPS O-antigen and outer core minimized unstructured OM density interfering with GBP1 resolution at the inner leaflet (38). Farnesylated GBP1 purified directly from human cells as a single FPLC peak harboring the lipidated species also ensured proper OM membrane anchorage and insertion. These modifications combined to yield the first sub-tomogram averaging of *Stm*-bound GBP1. Further efforts should help refine the extended GBP1 conformer and its bidomain configuration below ∼30Å in the future.

Together, our 3D reconstruction from cryo-ET elucidates the mesoscale architecture of a unique host defense structure - the GBP1 coatomer - inside the human cytosol. Our findings reinforce the importance of higher-order protein assembles within the innate immune systems of animals and plants; these nanomachines concentrate signaling and effector proteins for rapid mobilization of cell-autonomous and systemic resistance to infection. Our work also revealed that bacteria perceive encapsulation by this supramolecular complex and tune their responses accordingly, in this case, by regulating periplasmic LPS release that trigger caspase-4 activation directly atop the coated bacterium. Such crosstalk highlights the ongoing host-pathogen arms race and how IFN-*γ*-induced GBPs can overcome microbial evasion strategies to help defend the host cell interior.

## Acknowledgements

We thank J. Nikolaus, A. Tunaru, G. Torrence, E. Groisman and X. Liu for experimental advice and technical help.

## Author contributions

S.X., C.J.B, A.M. E-S.P., B-H.K., and J.D.M. conceptualized the project and helped write the paper. S.X., C.J.B, A.M. E-S.P., B-H.K., P.K., S.H., and Y.Z. performed all experiments with external collaboration by J.B. All authors discussed the results and commented on the manuscript.

## Data Availability

All data are available in the main text or the supplementary materials.

## Supplementary Materials

### Antibodies & Reagents

#### Antibodies

Antibodies used were anti-Flag M2 (Sigma), anti-HA (16B12 Biolegend), anti-Myc (9E10; ThermoFisher), anti-GFP (11814460001; Roche), anti-GBP1 (sc-53857), anti-GBP2 (1E5; Origene), anti-*Salmonella* O Group B antiserum (BD), anti-flagellin (FliC-1; BioLegend), anti-*β*-actin (ab6276), anti-GAPDH 41335; SCBT), anti-IL-18 (PM014; MBL) anti-Caspase-4 (clone 4B9; Enzo), anti-GSDMD (64-Y; SCBT).

#### Reagents

Recombinant *Podisus maculiventris* thanatin (*E. coli-*derived, MyBiosource), Murepavadin TFA (POL7080; MedChemExpress), guanosine 5′-triphosphate sodium salt, GTP-*γ*-S (tetralithium salt), guanosine 5′-diphosphate sodium salt (GDP), aluminum trifluoride, chloramphenicol, S-(3,4-dichlorobenzyl)isothiourea (A22, hydrochloride), Ficoll and biotin (Sigma); recombinant human IFN-γ (285-IF/CF; R & D Systems).

### Bacterial strains

Bacterial strains were generated in-house or kindly provided by the following groups: *Salmonella enterica* serovar Typhimurium (*Stm*) strain 1344 and flagellin-deficient *Stm* ^Δ*flhD*^ (Dr. Jorge Galan); (*48*) *Stm* UK-1 wildtype, Δ*wzy*, Δ*waaL*, Δ*waaJ*, Δ*waaI*, Δ*waaG*, Δ*lpxR*, Δ*pagL*, Δ*pagP*, and χ11088 (*Stm^ΔlpxRΔpagLpagP^* triple mutant) (Dr. Roy Curtiss III) (27, 29); *Pseudomonas aeruginosa* L2 strain (Dr. Barbara Kazmierczak), *Bacillus subtilis* (Jun Liu) and *Listeria monocytogenes* 140203S (Dr. Herve Agaisse).

The following 15 Stm strains were constructed in-house on an 1344 isogenic background: _StmΔminD, StmpBAD::ftsZ, StmmreB(K27E), StmmreB(D78V), StmHA-rseA::mCherry, StmlpxC-Flag::mCherry, StmPrpoE:sfgfp:::mCherry, StmPfkpA:sfgfp:::mCherry, StmtreA:sfgfp:::mCherry, StmPpmrD:sfgfp:::mCherry, StmPpagP:sfgfp:::mCherry, StmmScarlet-I, StmeGFP, StmmRFP and StmAIDA-C-BirA. In addition, StmΔWaaG::pBAD-ftsZ_ was generated on the UK-1 background for cryo-ET. A complete list of available strains and their construction will be available at https://medicine.yale.edu/profile/john_macmicking/.

### Bacterial infections, LDH assay and IL-18 ELISA

For *Stm* infections, overnight bacterial cultures were diluted 1:33 in fresh LB, grown for 3h before being washed once in PBS and used to infect HeLa cells at 80% confluence with an MOI of 5-20 as indicated. Plates were centrifuged for 10 min at 1000 x *g* and incubated for 30 min at 37°C to allow invasion. Extracellular bacteria were killed by replacing media with fresh DMEM containing 100 μg/ml gentamicin for 30 min. Cells were washed 3 times and incubated with 20 μg/ml gentamicin for the duration. To enumerate live bacteria, cells were lysed in PBS + 0.5% Triton X-100 and serial dilutions plated on LB agar. For LDH assay

### CRISPR-Cas9 cell lines & stable complementation

To generate stable gene knockouts in HeLa CCL2 cells, sgRNAs were cloned into pX459 (Addgene) per established protocols. 2-4 sgRNAs targeting each gene (200 ng total DNA) were transfected in 24 well plates for 24 h, followed by selection with 1 μg/ml puromycin for 48 h. Surviving cells were expanded into media lacking puromycin for 48 h, then subject to limiting dilution to obtain single colonies. Colonies were screened first by PCR, then by western blot and the genotype of each positive clone determined by Sanger sequencing. The following CRISPR-Cas9 mutants were generated: GBP1^-^/^-^, GBP2^-^/^-^, GBP3^-^/^-^, GBP4^-^/^-^, GBP1^-^/^-^2^-^/^-^ double mutant, GBP*^Δ^*^1q22.2^ septuple mutant (GBP1^-^/^-^GBP2^-^/^-^GBP3^-^/^-^GBP4^-^/^-^GBP5^-^/^-^GBP6^-^/^-^GBP7^-^/^-^), CASP4^-^/^-^, GSDMD^-^/^-^, and AHOH^-^/^-^.

In addition, we generated a series of cell lines stably or transiently complemented with GBP mutants, affinity probes or reporters. These included GBP1^-^/^-^ clonal lines complemented with either of the following: EGFP-GBP1, RFP-GBP1, mNG-GBP1, EGFP-GBP1^S52N^, EGFP-GBP1^DD103,108NN^, EGFP-GBP1^D184N^, EGFP-GBP1^C589S^, EGFP-GBP1^C589S^, EGFP-GBP1*Δ^α^*^13ARR^, EGFP-GBP1*Δ^α^*^13RAR^, EGFP-GBP1*Δ^α^*^13RRA^, EGFP-GBP1*Δ^α^*^13ARA^, EGFP-GBP1*Δ^α^*^13AAR^, EGFP-GBP1*Δ^α^*^13AAA^, BioID, BioID-GBP1, and BioID-GBP1 ^C589S^. Here alanine scanning mutations in the C-terminal polybasic patch (a.a. 584-586) of GBP1 were introduced according to Stratagene Quickchange protocol and confirmed by DNA sequencing.

We also complemented CASP4^-^/^-^ with GFP-caspase-4 and GSDMD^-^/^-^ with either GFP-FL-GSDM GFP-NT-GSDMD, or GFP-CT-GSDMD. A complete list of available knockouts and their construction will be available at https://medicine.yale.edu/profile/john_macmicking/.

### Cell culture and transfection

HeLa (CCL2) and 293T cells were purchased from the American Type Culture Collection (ATCC). U2OS cells were a gift from Grant Jensen (Caltech). Cells were grown in DMEM supplemented with 10% (v/v) heat-inactivated fetal bovine serum (FBS) at 37°C in a 5% CO_2_ incubator. Lentiviral (LentiCrisprV2) or retroviral (pMSCV-puro) transductions were done by incubating dilutions of 0.45 μm filtered supernatants from transfected 293T cells with 8 μg/ml polybrene for 24 h. For selection of stable transductants, 1 μg/ml puromycin was included. For transient transfections, *Trans*IT^®^-LT1 (MIRUS) was used according to manufacturer’s instructions. To minimize toxicity in microscopy experiments, 200 ng of DNA was transfected per 24 well cover slip. When required, HeLa cells were stimulated with 500-1,000 U/ml IFN-*γ* for 18 hours.

For infections with *Stm* expressing arabinose-inducible LpxC-Flag or HA-RseA, SPI-1-induced bacteria were first treated for 3h with 0.4% L-(+)-arabinose to induce HA-RseA expression in axenic culture. IFN-*γ*-activated HeLa cells of different genotypes were infected with these bacteria at an MOI of 20 and incubated for 30 min to allow invasion. Culture medium was replaced with new medium containing gentamicin (100ug/ml) and incubated for 45 min. A second medium replacement followed with gentamicin (10ug/ml) for 1 hr. Cells were lysed in 0.5% Triton X-100 and low speed centrifuged (4,000 x g) to pellet cell debris and vacuolar organelles before collecting the cytosolic fraction at 14,000 x g. Cytosolic lysates were immunoblotted with anti-HA or anti-Flag antibodies. mCherry expression from a constitutive ribosomal *rpsM* promoter was immunoblotted as an internal loading control loading for highly expressed RseA. A single non-specific band (nsb) of high molecular weight detected by the same anti-Flag antibody as LpxC-Flag was chosen to serve as a robust internal control because LpxC was detected at very low levels within human cells. For infections with *Stm* 1344 reporter strains (*pFCcGi::PrpoE-sfgfp, pFCcGi::PfkpA-sfgfp, pFCcGi::PtreA-sfgfp, pFCcGi::PpagP-sfgfp,* and *pFCcGi::PpmrD-sfgfp* co-expressing mCherry), bacteria were introduced at an MOI of 20 and examined at 3 hours post-infection. Both IFN-*γ*-activated cells and unactivated GBP1*^Δ^*^1q22.2^ cells reconstituted with IFP-GBP1 were tested for analysis. The latter group established sufficiency and avoided other IFN-induced proteins causing alterations in reporter expression (eg. APOL3; 12). Mean fluorescence intensity (MFI) for each channel enabled GFP:RFP expression ratios to be established across multiple regions of interest (ROIs).

### CLICK chemistry & metabolic labeling

#### FPP-azide-biotin CLICK chemistry

293E cells expressing GFP or GFP-GBP1 mutant constructs were plated at 2×10^5^ cells in each well of a 6-well plate. Cells were then treated with 20uM Azido farnesyl pyrophosphate (C10248) for 18 hours followed by cell lysis in 500uL of 20mM Tris(7.5) 100mM NaCl 1%TX-100 buffer containing Roche protease inhibitors. Cells were further sonicated (30% power, 3 times for 1 minute each) on ice to liberate membrane bound proteins. Lysates were centrifuged at 21000g for 10 minutes. Supernatants were transferred to a new Eppendorf tube and were treated with 100uM Biotin DIBO (C10412) overnight at room temperature in the dark. 1ug Roche monoclonal anti-GFP antibody was added to each IP followed by a 2-hour incubation at 4°C. 20uL of pre-equilibrated Protein G Sepharose (17-0756-01) was added to each reaction for an additiona 2 hours. Beads were pelleted at 4000g for 5 minutes followed by 8 washes in lysis buffer. Samples were subsequently eluted by 100uL addition of 2X SDS-Sample Buffer followed by heating at 100°C for 20 minutes for immunoblotting with Immunoblots were performed with streptavidin-HRP and Roche anti-GFP.

#### KDO-azide Cu^2+^-free CLICK chemistry

The 3-Deoxy-D-manno-octulosonic acid (KDO) of log-phase *Stm* was labelled using CLICK-mediated according to the manufacturer’s instructions (Jena Bioscience). Briefly, an azide modification of the C8-position of KDO with a biorthogonal azido group is introduced which prevent reverse metabolism by KDO-8-P phosphatase. This 8-azido-8-deoxy-KDO modification enabled a biotin group within a dibenzocyclooctynol (DIBO) alkyne dye intermediate to be introduced via Cu(I)-free CLICK chemistry. For infection experiments, we conducted click chemistry reactions on bacteria that had already incorporated fluorescently labelled D-alanine into the underlying peptidoglycan scaffold. Unincorporated DIBO was removed via extensive washing in PBS.

#### Metabolic labeling of Stm peptidoglycan & LPS release in situ

Fluorescent blue or red D-alanine analogs (HCC-amino-D-alanine, HADA; TAMRA-amino-D-alanine) was incorporated as described (33). Infection of HeLa cells at MOI 20. After 40 minutes of infection the media was changed to DMEM with 100 ug /ml gentamycin, after next 1 hour to 30 ug/ml gentamycin. Cells were fixed with PFA at 2 hours post infection and immune-stained for GBP1 or LPS. At least 10 GBP1-positive and 10 GBP1-negative fields of view were collected using OMX Blaze SIM microscope (GE) in the SIM mode. All images were subjected to processing to widefield image, de-convolution and maximum intensity projection for semi-automatic analysis in CellProfiler. Detection of objects was based on sum of all channels (sum of area of all channels = 100%, HADA/TAMRA, GBP1, LPS – object detection) and signal area was measured.

### Microscopy

#### OMX-SR Blaze & Multicolor Confocal Microscopy

HeLa cells were seeded on 12 mM high performance cover glass #1.5h (Thorlabs). For live imaging, cells were seeded on 4-well chambers with 1.5 high performance cover glass (Cellvis). Cells were seeded 48 h prior to imaging to reach 80% confluency on the day of infection and treated with 500 U/ml IFN-*γ* where required for 18-24 h prior to imaging. To image bacterial infections, bacteria were added to cells as described for infections at an MOI of 20. Images were analyzed on a DeltaVision^TM^ OMX SR Blaze microscopy system (GE Healthcare) or a laser scanning confocal model SP8 (Leica).

For ultrafast live imaging, GBP1^-^/^-^ HeLa cells were transfected with RFP-GBP1, induced with 1000 U/ml IFN-γ for 18 hr and infected with EGFP-*S.tm* at MOI 20. After 40 minutes, the media was changed to DMEM with 30 ug/ml gentamycin and the sample was imaged starting 60 minutes post-infection at 37C, 5% CO_2_. Images were collected every 45 s using OMX-SR Blaze microscope (GE) in 1024 x 1024-pixel mode at ∼65 frames.sec^-1^. Images represent max. intensity projections of deconvolved z-stacks (Softworx, GE). Post-acquisition calculations for real-time voxel (boxed) assembly events used Imaris (Oxford instruments) software. When combined with atomic structure volumes of GBP1 (PDB 1F5N) and RFP (PDB 1GGX) the total number of spatially constrained RFP-GBP1 molecules per voxel were counted, along with coatomer kinetics. Long-term imaging used low light OMX-SR conventional mode for up to 3 hours continuous recording at 5-minute intervals.

For examining the COATOMER_450-708_ complex, we constructed fluorescent fusion proteins for GBPS1-4, caspase-4 and GSDMD across different spectral range to accommodate 5 or possibly 6 proteins simultaneously along with *Stm*. A color-coded matrix of 48 proteins was generated by fusing each of the 6 coatomer proteins to each of the following fluorescent reporters: mAzurite, mSapphire, pmTurqouise2, pmEmerald, pmVenus, pmOrange, pmCardinal, pmIFP24 (Addgene); E_x_/E_m_ range, 384/450-684/708nm. Multi-arrayed testing found 5 combinations of either pmIFP24-GBP1, pmOrange-GBP2, pmVenus-GBP3, pmEmerald-GBP4, Sapphire-caspase 4 or -GSDMD, or pmOrange-GSDMD could be used successfully with *Stm*. COATOMER_450-708_ combinations were transfected into the GBP*^Δ^*^1q22.2^ septuple mutant, activated with IFN-*γ* and infected with *Stm*-Alexa568 for localization. Five or 6-color images were captured on a Leica SP8 confocal microscope using a series of lasers to excite fluorescence across this broad wavelength and avoid bleed-through as shown in fig. S2D. The corresponding laser parameters (laser, % power, wavelength reception band) used were: pSapphire (Diode 405, 4, 424-444), pmEmerald (WLL 470, 20; 490-520), pmVenus (WLL 516, 6; 535-559), pmOrange (DPSS 561, 25, 566-593), pmIFP24 (HeNe 633, 75, 680-730).

### Whole-cell 4Pi single-molecule switching nanoscopy (W-4PiSMSN)

For W-4PiSMSN imaging, a customized microscope built using a vertical 4Pi cavity around two opposing high-NA objective lenses as detailed elsewhere (17) was used to capture high-resolution images of the GBP coatomer in IFN-*γ*-activated Hela cells. Cell samples were prepared on 25 mm diameter round precision glass cover slips (Bioscience Tools, San Diego, CA) that had been immersed in 1M KOH and sonicated for 15 min in an ultrasonic cleaner (2510 Branson, Richmond, VA). Sequential washes in Milli-Q water (EMD Millipore, Billerica, MA) and sterilization with with 70% ethanol was followed drying and poly-lysine coating of coverslips. HeLa cells were grown on coverslips for 24–48 hour before fixation in 4% paraformaldehyde. Anti-GBP1 (1B1; Santa Cruz; 1:500 dilution) and GBP2 (1E5; Origene; 1:200 dilution) antibodies were detected by AF-647 and Cy3B secondary antibodies as described (17). Placement of fixed samples on coverslips were held in the sample frame using an additional curing silicone and imaged immediately after the silicone solidified. Subsequent W-4PiSMSN data acquisition enlisted four phase images that were arranged along the splitting line of the sCMOS camera’s upper and lower readout region. 50,000 to 320,000 camera frames were recorded at 50 or 100 fps. These phase images were merged into a single image using a transformation matrix obtained from a combination of algorithms using log-polar and affine transformations (17). Elliptical Guassian models to detect sCMOS camera noise for elimination enabled single molecule positions to be obtained along photons, background and log likelihood ratios. For two-color images of endogenous GBP1 and GBP2, phase shifts between *s* and *p* polarization for the two-color channels differed by 0.3 radians. Drift correction further aided positioning of emitted photons to yield W-4PiSMSN images.

### Purification of recombinant proteins

*Guanylate binding proteins 1-4 (GBPs1-4) and their mutants, caspase-4^C258A^, RFP-AtGBPL1* The coding sequences of human GBP1 and its mutants were cloned into a customized vector pCMV-His_10_-Halo-HRV-mRFP-TEV for coating assays. HEK293f suspension cells (a gift from Dr. James Rothman; mycoplasma-negative) was maintained at a concentration of 0.4×10^6^∼4×10^6^ cells/ml in Expi293 expression medium (ThermoFisher Scientific). 24 h prior to transfection, cells were seeded at a concentration of 1.2 ×10^6^ cells/ml. For transfection, cells were harvested and resuspended in fresh medium at a concentration of 2.5 ×10^6^ cells/ml. Cells were transfected by adding pCMV-His_10_-Halo-HRV-mRFP-TEV-containing clones to a final concentration of 1 µg/ml in media containing PEI at a concentration of 5 µg/ml. 24 h after transfection, cells were diluted 1:1 (v/v) with fresh medium containing 4 mM valproic acid and cultured for an additional two days. 2 x 10^9^ cells were harvested via centrifugation (500 x *g*, 10 min), washed once in cold PBS, resuspended in lysis buffer (50 mM HEPES, pH7.5, 500 mM NaCl, 1 mM MgCl_2_, 10% glycerol, 0.5% CHAPS, 1 mM TCEP) and lysed via sonication. Cells were cleared at 35,000 x *g* for 1 h at 4°C. Supernatant was collected and incubated with 1 ml bed volume of HaloLink resin (Promega) at 4°C overnight with gentle rotation. The resin was sequentially washed twice (10 min each) with wash buffer 1 (50 mM HEPES, pH7.5, 500 mM NaCl, 1 mM MgCl_2_, 10% glycerol, 0.5% CHAPS), wash buffer 2 (50 mM HEPES, pH7.5, 1 M NaCl, 10% glycerol) followed by wash buffer 1.

To elute bound proteins, Halo resin was resuspended in lysis buffer and digested with homemade GST-HRV-His protease overnight at 4°C with gentle rotation. Resin was pelleted and the HRV protease was removed from the supernatant via Ni-NTA beads by affinity chromatography (QIAGEN). Flow through was collected, concentrated and further purified and buffer-exchanged via size exclusion chromatography (Superdex® 200 Increase; GE Healthcare) equilibrated with storage buffer (20 mM HEPES [pH 7.5], 150 mM NaCl, 1 mM MgCl_2_, 1 mM TCEP). Fractions were analyzed by SDS-PAGE, pooled, concentrated and flash frozen in liquid nitrogen before storing at −80°C. Protein concentration of rRFP-tev-hGBP1 (rRFP-hGBP1 for short) was determined via BCA and SDS-PAGE electrophoresis with BSA standards running aside.

Recombinant human GBP1, GBP2, GBP3, and GBP4 as well as the GBP1 mutants (hGBP^1S52N^, hGBP1^D184N^, hGBP1^DD103,108LL^, hGBP1^C589S^ and hGBP1^R3A^) were produced by cloning the respective genes into the mammalian expression vector, pCMV-3Tag −1, with an N-terminal FLAG tag. Human caspase-4 ^C258A^ was cloned into pcDNA 3.1/Hygro for N-terminal FLAG attachment. Plasmids were transfected via Mirus LT1 into HEK-293 E cells. Cells were harvested after 14 h and lysed for 2 h in buffer 50 mM Tris, 150 mM NaCl, 5 mM MgCl_2_, 1 mM DTT containing 1% Triton X-100 and protease inhibitor. Supernatant containing the expressed protein passed through anti-Flag M2 affinity gel and bound Flag protein eluted using Flag peptide 150 ug/mL in buffer 50 mM Tris, 150 mM NaCl, 5 mM MgCl_2_ and 1 mM DTT. Further chromatographic purification was undertaken on an AKTA FPLC (GEAmersham) instrument. All recombinant proteins made in human cells were subject to Limulus amebocyte assay (LAL; ThermoFisher) to confirm the absence of LPS contamination (<0.01 EU/mL, lower detection limit).

### Thin-layer Chromatography

Thin layer chromatography (TLC) was used to separate GTP, GDP and GMP and performed exactly as previously described (7–9). Briefly, α-[^32^P]GTP (Perkin Elmer) hydrolysis by purified recombinant proteins in reaction buffer (20 mM HEPES [pH 7.0], 150 mM NaCl, 5 mM KCl, 1 mM MgCl_2_, 100 μM GTP, 10 μCi α-[^32^P]GTP) was determined at 25°C before quenching the reaction with 142 mM EDTA after 7 h. The resulting products were separated by PEI cellulose (Sigma, Cat#Z122882) with fluorescent indicator (UV 254) using 750 mM KH_2_PO_4_ (pH 3.5) as solvent and visualized by autoradiography.

### Limulus Amebocyte Assay (LAL) for lipid A detection

Overnight-cultured *Stm* was pre-incubated with indicated inhibitors at 37°C with shaking for 2h. The Lpt inhibitors included thanatin (10ug/ml) or murepavadan (0.05ug/ml). After centrifugation by 3,500 rpm room temperature for 20min, bacteria were washed three times by 10-fold volume of coating buffer (50mM HEPES (pH7.5), 150mM NaCl, 1mM MgCl_2_ and 1mM DTT) and finally resuspended by 1ml of coating buffer containing 2mM GTP or GTP*γ*S. Bacteria aliquoted were incubated with or without 4μM of recombinant GBP1 or its mutants at 30°C for 1 hour. Supernatants were carefully collected by centrifugation (8,000 rpm, RT for 2 minutes) and diluted 1:1000-1:5000. Released LPS was measured by ToxinSensor™ Chromogenic LAL Endotoxin Assay Kit (GenScript) according to manufacturer’s instructions.

### LPS-binding Assays

Human GBP1, GBP2, GBP3, GBP4, GBP1 mutants (GBP1^S52N^, GBP1^D184N^, GBP1^DD103,108LL^, GBP1^R53A^ and GBP1^C589S^) and caspase-4^C258A^ proteins were expressed as Flag-tagged proteins in human embryonic kidney (HEK)-293E cells and isolated from large-scale adherent cell culture using Flag M2 beads. Recombinant human proteins were further purified to single peak via FPLC. Fluorescence anisotropy assays were conducted at 37°C across different GBP concentrations in binding buffer (50 mM Tris, 150 mM NaCl, 5 mM MgCl_2_, 0.3 mM GDP and AlF_X_ [10 mM NaF, 0.3 mM AlCl_3_], pH 7.0) and Flag-caspase-4^C258A^ (50 mM Tris, 150 mM NaCl, 1 mM DTT, pH 7.0) for 15 min followed by addition of *Salmonella* minnesota LPS-Alexa Fluor 488 (ThermoFisher) to a final concentration 250 nM. After 1 hour incubation with LPS, fluorescent anistropy values measured by SpectraMax i3x (Molecular Devices).

### Reconstituted Coatomer Assays

Wild-type *Stm* and *Stm* mutants were freshly streaked on LB plates. A single fresh colony of bacteria was cultured in LB medium overnight at 37°C. Before coating assay, bacteria were diluted 1/60 in fresh LB medium, cultured for additional two hours and harvested via centrifugation (4000 g, 5 min at RT). To remove excessive free LPS which inhibits rRFP-hGBP1 targeting, bacteria was washed twice in coating buffer (50 mM HEPES [pH7.5], 150 mM NaCl, 1 mM MgCl_2_) and further diluted to an optical density (OD_600_) of 0.1 in coating buffer. Immediately before coating, rRFP-hGBP1 was thawed at room temperature and centrifuged at 12,000 g for 15 min to remove any insoluble aggregates. For coating, 2 *µ*M rRFP-hGBP1 and 2 mM GTP were added to each 0.1 OD of bacteria, gently mixed and incubated for 1 hour at RT before imaging.

To image rRFP-hGBP1-coated bacteria, coating reactions (20 *µ*L) were transferred to 384-well glass bottom plate (Cellvis; cat#P384-1.5H-N). The plate was centrifuged at 2000 g for 1 min to collect all liquid to the bottom of the well. All images were acquired using a Leica SP8 laser-scanning confocal microscope. Briefly, analyses were performed with a 63×/1.40 oil immersion objective. Focal plane was set to the bottom of the well. RFP was excited at 561 nm and detected at 590−610 nm. The optical slices were acquired in confocal mode (1 Airy unit) with an average of six scans. Images were collected in a 512×512 format. Image analysis was performed with FIJI/ImageJ.

For coatomer assembly, samples were doubly diluted from 2μM rRFP-GBP1 until loss of coating to establish a coating curve. The observed behavior was plotted with best-fit interpolation and a sigmoidal curve emerged. Curve fitting together with regression analysis found half maximal and Hill slope values using Graph Pad Prism 9.1.1.

### In vitro Phase Separation

Protein aliquots of rRFP-GBP1 and rRFP-GBPL1 (7) were thawed at room temperature, centrifuged at 14,000 × g for 5 min to remove any aggregated protein. Droplet formation was induced by diluting protein to low salt buffers (50 mM HEPES, pH7.5, 150 mM NaCl, 1 mM MgCl_2_) by mixing with various volumes of no salt buffer (20 mM HEPES [pH 7.5], 1 mM TCEP) and analyzed in chambered coverglass (Grace Bio-labs) using a Lecia SP8 laser scanning confocal at 63×/1.40 magnification. Addition of Ficoll at 5%, 10%, 15%, 20% w/v had no effect on rRFP-GBP1.

### Electron Microscopy

#### Negative-stain electron microscopy

To visualize rRFP-GBP1 on minicells, 100µL minicells were washed and dissolved in the HEPES buffer (pH 7.4) to the OD_600_ equal to 0.1. The minicell suspensions were incubated with rRFP-GBP1 with a final concentration of 2 µM in the presence/absence of GTP at room temperature for 1 hour. The mixture was concentrated to 10µL for further TEM observation. 5µL RFP-GBP1 coated and non-coated minicells were loaded onto glow-discharged carbon film coated cooper EM grids (EMS, cat#CF400-Cu-50) for standing 1 minute. The EM grid was washed and stained with 2% uranyl formate for 30 seconds. The grid was blotted by filter paper (Whatman FILTER PAPERS 2 cat No. 1002-090) and dried for 1 minute. Grids were examined in JEOL1400 plus electron microscope with acceleration voltage of 80 kV.

### Cryo-EM Sample Screen

Before the cryo-ET data collection of the minicell sample, EM grids were transferred into 200 kV cryogenic electron microscopy (Thermo Fisher Glacios) equipped with a Gatan K2 camera. Images were taken at eucentric height with a defocus range from −3.5 µm to −4.5 µm. The images were collected at the pixel size of 1.478Å with an overall dose of 40 e-/Å^2^. The drifts in the subframes were corrected by Motioncorr package. Corrected screening images were visualized by IMOD package.

### Cryo-electron Tomography Cell Seeding on EM Grids

Pre-treatment of EM grids. EM grids (Quantifoil R1/4 gold 200 mesh, Cat#Q250AR-14) were placed onto 18 mm^2^ cover-glass that had been washed and stored in 100% ethanol. They were then placed into glass bottom dishes (Cellvis, 35 mm dish with 20 mm micro-well cover glass) and treated with 70% ethanol under UV illumination for 10 min at room temperature^52^. EM grids were washed 6 times with sterile and degassed water before treatment with 0.05 mg/mL collagen I (Gibco, cat, A10644-01) for 60 min in a 37°C incubator followed by washing in degassed/distilled water. Collagen-coated grids were incubated in DMEM medium overnight at 37°C incubator with 5% CO_2_ for the further use.

The GBP*^Δ^*^1q22.2^ septuple mutant was cultured in 10 cm dish to the density of 4,000,000 cells per dish. The HeLa cells were transfected with a plasmid encoding hGBP1 fused with GFP at the N terminus using Mirus LT1 kit. In next day, cells were washed by PBS and treated with 0.05% Trypsin-EDTA for the detachment. The detached HeLa cells were centrifugated and resuspended with DPBS buffer. The HeLa cells were filtered by a 40 µm nylon mesh filter (Fisher Scientific, cat no. 22363547) to remove the junks. The filtered HeLa cell suspension were transferred into 5mL polystyrene round-bottom tube (Corning, REF, 352235) with cell-strainer cap for the further FACS sorting process. The HeLa cells with GFP signals were sorted into the collection tubes. The cells were centrifugated and resuspend to a final cell density of 20,000 cells/mL in DMEM supplemented with 10% fetal bovine serum and 100 µg/mL Pen Strep (Gibco, REF, 15140-122), seeded on the pre-treated EM grids and further incubated for overnight. For the infection assay, *Stm* bacterial cells transformed with a plasmid encoding salmonella FtsZ induced by 0.2% L-arabinose were cultured to a density of OD_600_ equal to 1. The HeLa cells on the EM grid were infected by elongated Stm1344 at MOI 200. The EM grids coated with HeLa cells and bacterial cells have a further centrifugation at 800g for 5 minutes for spinning the bacterial cells down to HeLa cell surface. The EM grids were transferred into 37°C incubator for additional 20 minutes. The EM grids were washed by DPBS for 3 times and incubated with 100 µg/mL gentamycin contained DMEM medium in the supplement of 10% FBS for additional 1 hour infection.

For the minicell sample preparation from *Stm*^ΔminD^, bacterial cultures were grown overnight at 37°C in LB medium in the presence of 100 µg/mL ampicillin. 10 mL bacterial overnight cultures were added into 1 L fresh LB medium in the presence of ampicillin and were grown to late log phase. The whole cultures had a 3-step centrifugation: the cultures were centrifuged at 6,240 g (Beckman coulter, Avanti JXN-26, rotor, JLA-8.1000) for 10 min to remove the normal bacterial cell; the supernatant was centrifuged at 24,820 g (rotor, JLA-16.250) for 10 min to collect minicell. The minicell fractions were suspended by HEPES buffer (pH 7.4) and filtered by 0.45 µm PVDF membrane (Merck Millipore, REF, SLJVM33RS). The fraction was centrifuged at 20,000 g (rotor, JA-25.50) for further 10 min. the minicell fraction was adjusted to OD_600_ equal to 1 for further rRFP-GBP1 in vitro coating experiment.

For the minicell sample preparation from FtsZ overexpressed ΔwaaG strain, a fresh second culture was grown in LB medium containing 100 µg/mL ampicillin and 0.2% L-arabinose for a continuous FtsZ induction for 20 hours in the 37°C. The minicells collection from ΔwaaG adopted the same protocol.

### Vitrification and Cryo-correlative Fluorescence Light Microscopy (CLEM)

EM grids seeded with HeLa CCL2 cells expressing EGFP-GBP1 were pre-screened using a fluorescent light microscopy, clipped on a custom manual plunger, blotted using Whatman #1 filter paper (GE, cat#1001-110), and frozen in liquid ethane by the plunger^53^. Frozen grids were transferred onto a cryo-stage and clipped within the O-ring and C-ring (ThermoFisher Scientific, cryo-FIB autogrid and C-clip, respectively). Cryo-CLEM (Leica Microsystems EM Cryo-CLEM microscope) was performed as previously described^54^. Grids were transferred into Leica-CLEM cartridge docked at a pre-cooled shuttle docking station and then transferred onto a cryo-stage equipped with a pre-cooled 40× objective lens. A spiracle 4×4 grid region was selected, and 16 sub-regions were imaged. Montage GFP-positive images were acquired on the selected area using the Z-stack mode within a stepwise of 0.35 µm. This montage served as the navigating map in subsequent milling assays by the focused ion beam method.

### Milling Lamella by Focused Ion Beam (FIB)

Cryo-CLEM demarcated grids were transferred into cryo-DualBeam microscope equipped with cryo-stage and cryo-transfer shuttle systems (Thermo Fisher Scientific, Aquilos cryo-FIB focused ion beam/scanning electron microscope). The lamella was prepared according to Aquilos cryo-FIB protocols. The sample on the O-ring side of the grid was sputter-coated with platinum (1 kV, 30 mA, 10 pa, 15 s) to increase sample conductivity during FIB milling. MAPS software was used to capture an EM grid montage in the electron beam mode before importing the fluorescent montage from cryo-CLEM to generate a merged map to establish potential targets for milling lamella. Eucentric height of the targeting area was refined and stage tilting angle determined at 16° for lamella. At target position on the grid organometallic platinum coating was sprayed for 5 seconds by a gas injection system (Thermo Fisher Scientific, GIS). Initially, rough milling was starting from top-side with 300 pA current and continued in a small stepwise manner. Concurrently, SEM images were taken to observe intracellular elongated bacteria. Rough milling stopped on the top-side when the feature of elongated bacterial shape shows up. Rough milling on the bottom-side started with 300 pA current in a large stepwise manner. When the lamella thickness arrived to 1.5 μm distance, the milling current started to change to 100 pA until the thickness of the lamella is about 0.8 µm, the rough milling stopped. Fine milling was mainly focused on the bottom-side at < 50 pA. An electron beam was used to monitor the milling process at 5 keV and 13 pA. A final fine milling was done on the both side of the lamella. Upon completion of the milling, the grid was sputter-coated with platinum (15 mA, 10 Pa, 10 s) to increase conductivity of milled lamella. The grid was transferred out of the FIB machine using the transferring shuttle system and either stored in a grid box (SubAngstrom, Pin type grid box) in liquid nitrogen until further data collection or re-confirmed by CLEM for correct positioning of GBP1-coated bacteria.

### Cryo-electron Tomography and Image Processing

Tomography data for the FIB milled lamella sample were collected using a Titan Krios G2 transmission electron microscope (Thermo Fisher Scientific) equipped with a 300 kV field-emission gun, a Volta phase plate (phase shift located around 1/2 pi), a Quantum post column energy filter (20-eV slit) and a Gatan K2/K3 summit direct detection camera. The images were taken in a dose-fractionated mode at near focus using SerialEM software. The resulting physical resolution is 0.45 nm/pixel. A total dose of 80 e/Å^2^ distributed among 33 tilt images was undertaken covering angles from 39° to −57° at tilting step of 3° and starting the first tilt series at −9° in a continuous data collection mode. The dataset for the minicell were taken a dose symmetric scheme without using phase plate. The tilt series for minicells sample were taken at the physical resolution of 0.14 nm/pixel starting from 0° and covered angle ranges of 51° to −51° with a 3° increments, resulting in a total dose of 100 e/Å^2^.

Collected dose-fractioned data were subjected to motion correction for generating drift-corrected image stack files^59,60^. Stack files were aligned using patch-tracking function of IMOD. 3D tomograms were reconstructed from aligned stack files by SIRT (Simultaneous Iterative Reconstruction Technique) method using Tomo3D. For reconstruction of binned tomograms, the aligned tilt series was scaled to 3.6 nm as pixel size. IMOD was used for visualizing the tomogram.

Surface rendering of tomogram was done with EMAN2.23, and refined with UCSF chimera. Briefly, for ribosomes, a template-matching strategy was performed to determine all ribosome coordinates and orientations. A mammalian 80S ribosome structure (EMD-3418) determined by Volta phase plate cryo-ET, and bacterial ribosome cryo-EM structure (EMD-0076) (50) were low-pass filtered to 20 angstrom and scaled to 3.6 nm/voxel to match to the tomograms.

EMAN2.3 was employed to do segmentation on membrane, vesicles and hGBP1 coatomer features. The cryoEM map of type 3 secretion system (T3SS) needle complex from Salmonella typhimurium (EMD-11781) (51) was scaled to 3.6 nm/voxel and fitted into the position of T3SS in the tomograms.

### Sub-tomogram averaging

TomoSegMemTV and PySeg packages were used for sub-tomogram analysis. For initial membrane segmentation, 11 tomograms from ΔminD dataset and 27 tomograms from ΔwaaG with a binning factor of 8 were selected for membrane segmentation by using TomoSegMemTV (52). PySeg package was used for tracing and picking the particles located on the membrane based on Discrete Morse theory (53). The coordinates of picked particles were multiplied by 4 to obtain the particles coordinates information in the CTF corrected and weight back projection reconstructed tomograms within a binning factor of 2. Sub-tomograms were extracted and were initially aligned based on the feature of outer membrane by Relion package (54). After the outer membrane is well aligned, general classification function in the PySeg was performed to distinguish the membrane and non-membrane feature. The classes that showed the feature of membrane were merged together to have a further general classification by applying a sylinder mask on the top of the membrane. The classification results showing the extra density of GBP1-like feature on the top of the membrane were collected for further constrained refinement by Relion package. For the dataset from ΔminD, 9,843 particles were used and for the ΔwaaG/ftsZ dataset, 20,148 particles were included for sub-tomogram average refinement. Because hGBP1 coating is densely-packed, constraint refinement did not go further than 26 Å.

### In silico protein sequence analysis

For lipid-binding motifs in human GBP1 compared with other related GTPases, evolutionary tree history was inferred using the Maximum Likelihood method and JTT matrix-based model. A bootstrap consensus tree inferred from 10,000 replicates was taken to represent the evolutionary history of the taxa analyzed. Branches corresponding to partitions reproduced in < 50% bootstrap replicates were collapsed. Initial tree(s) for the heuristic search were obtained automatically by applying Neighbor-Join and BioNJ algorithms to a matrix of pairwise distances estimated using the JTT model, and then selecting the topology with superior log likelihood value. A discrete Gamma distribution was used to model evolutionary rate differences among sites (5 categories (+G, parameter = 2.8088)). Evolutionary analyses were conducted in MEGA X.

## Statistics

Data were analyzed by GraphPad Prism 9.1.1. software. Unless otherwise indicated, statistical significance was determined by t-test (two-tailed) or one-way ANOVA (Dunnett’s multiple comparison tests) with Holm-Sidak post-hoc test.

---

**Fig. S1.**
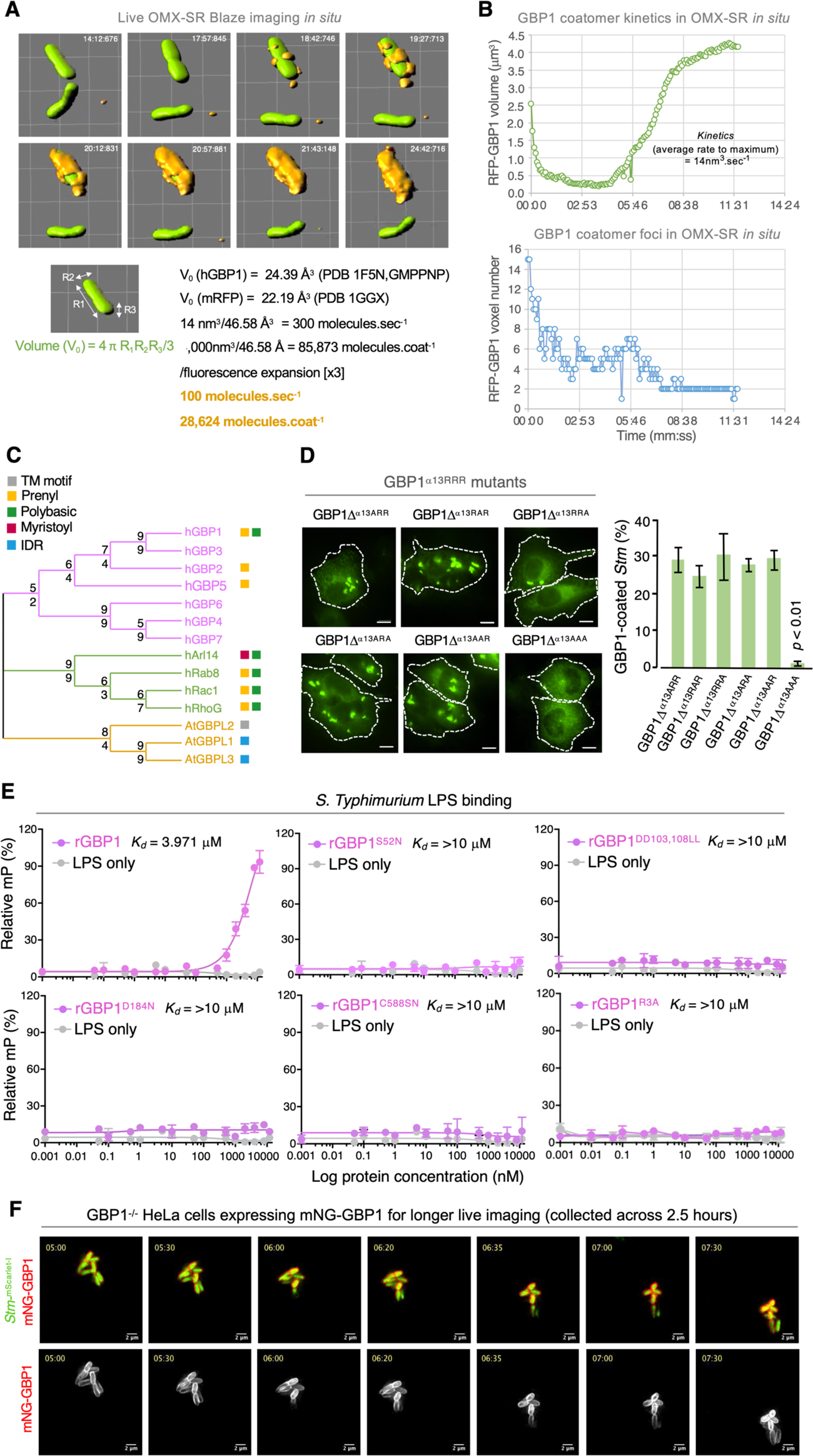
Human GBP1 coatomer characteristics and determinants of C-terminal anchorage. **(A)** Live OMX-SR Blaze imaging of EGFP-expressing *Stm* 1344 being coated by RFP-GBP1 at 2 hours post-infection. Volumetric and velocity measurements calculated by Imaris software (below) which incorporate crystal structure size and fluorescence enhancement. (**B**) The number of individual voxels diminished as the coat coalesced into a single polymeric platform. (**C**) Evolutionary tree of lipid-binding motifs in human GBP1 versus other related GTPases inferred via Maximum Likelihood methods and a JTT matrix-based model. Percentage of replicate trees where taxa clustered in 10,000 bootstrap replicates depicted next to branches. (**D**) Alanine scanning mutagenesis of the C-terminal polybasic patch (a.a. 584-586) of EGFP-fused mutants introduced into HeLa cells. Quantitation from more than 100 bacteria/group. (**E**) *Salmonella* minnesota LPS-Alexa Fluor 488 (ThermoFisher) binding curves to recombinant GBP1 and its mutants in fluorescence anisotropy assays. LPS alone did not contribute to polarization. Y-axis, % of maximal polarization (mP). Mean ± SD determined in triplicate for each protein concentration. Representative of 3-4 independent experiments. **(F)** Live widefield imaging of GBP1^-^/^-^ HeLa CCL2 cells transfected with mNeonGreen-GBP1, induced with 1000 U/ml IFN-γ for 18 hr and infected with mScarlet-i-*S.tm* at MOI 20. After 40 minutes of infection the media was changed to DMEM with 30 ug/ml gentamycin and the sample was imaged starting 300 minutes post infection. Images were collected every 5 minutes for the next 150 minutes and presented as maximum intensity projections of deconvolved z-stacks (Softworx, GE). Significance determined by one-way ANOVA with Holm-Sidak post-hoc test (D).

**Fig. S2.**
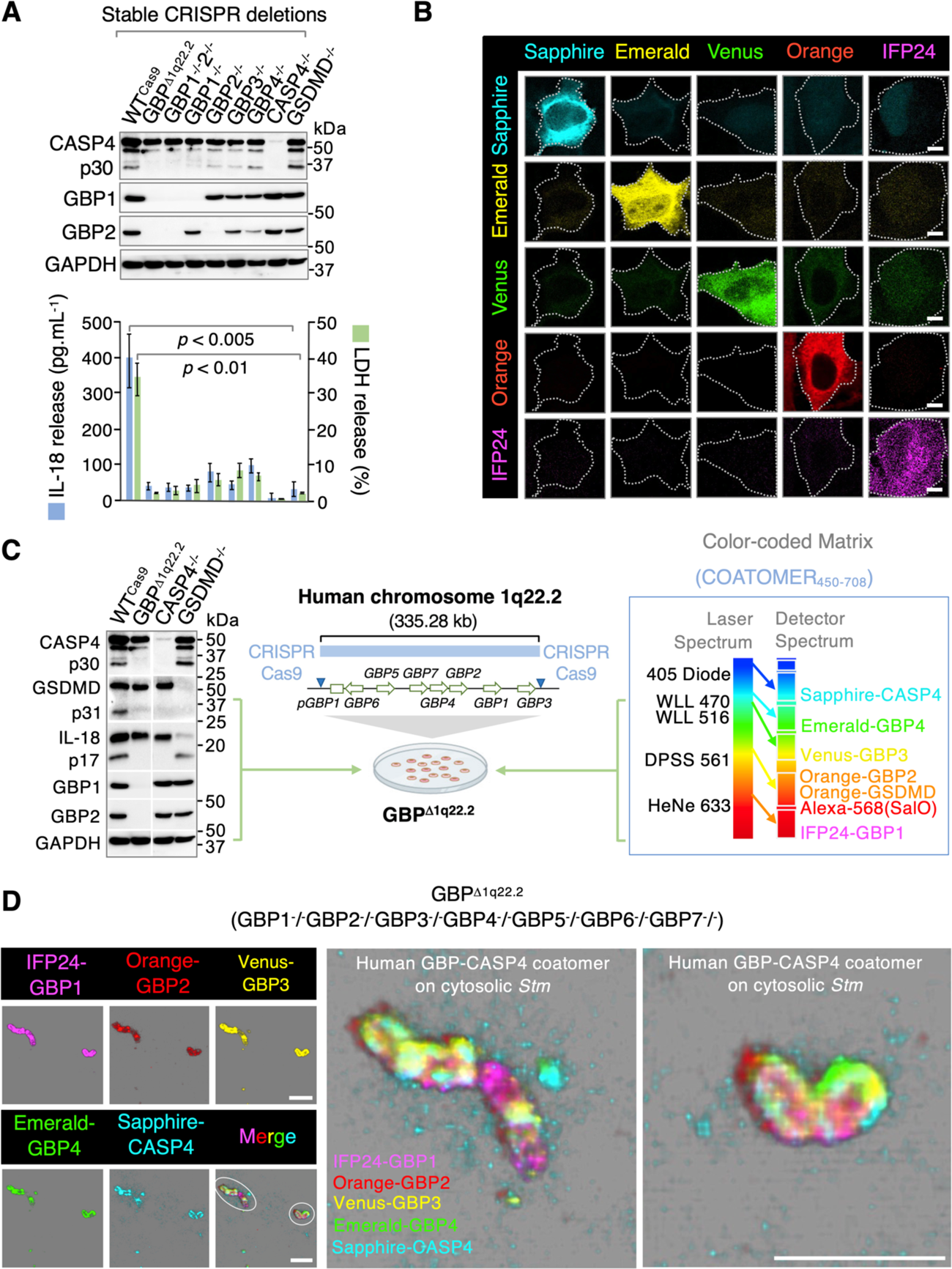
Functional coatomer partners and 6-member COATOMER_450-708_ reconstitution system. (**A**) (Top) Immunoblot of cleaved (p30) subunit of human caspase-4 in stably generated CRISPR/Cas9 HeLa cell lines activated with 500U/mL IFN-*γ* for 18 h before *Stm* infection. Endogenous GBP1, GBP2 and GAPDH shown as controls. (Bottom) IL-18 release detected by ELISA and LDH assay for pyroptotic cell death. Representative of 4-6 independent experiments performed in triplicate. Significance determined by one-way ANOVA with Holm-Sidak post-hoc test. (**B**) Establishing laser power and imaging conditions for multicolor confocal microscopy to reduce bleed-through from adjacent channels for 5-color imaging. HeLa cells simultaneously transfected with all 5 fluorescent protein plasmids used for tagging coatomer proteins and imaged across the 424-730 wavelength detector spectrum. Scale bar, 2μm. (**C**) (Left) Immunoblot of cleaved GSDMD p31 N-terminal fragment and cleaved p17 IL-18 fragment for phenotypic comparison of GBP*^Δ^*^1q22.2^ versus CASP4^-^/^-^ and GSDMD^-^/^-^ cells. Caspase-4 p30, GBP1, GBP2 and GAPDH included from (C) above as well for the same samples. (Center, Right) Introduction of the COATOMER_450-708_ plasmid array into GBP*^Δ^*^1q22.2^ cells depicting the removal of the complete human *GBP* gene cluster arrayed on chromosome 1q22.2 reconstitutes the coatomer. (**D**) Multi-color confocal imaging of the coatomer assembling on the same cytosolic bacilli in IFN-*γ*-activated GBP*^Δ^*^1q22.2^ HeLa cells. (Left) Individual channels with convergent assembly denoted by dashed circles. Scale bar, 2μm. (Right) Enlargement of the reconstituted coatomer in dashed circles depicting sub-regions occupied by different coat proteins on the bacteria surface. Scale bar, 2μm.

**Fig. S3.**
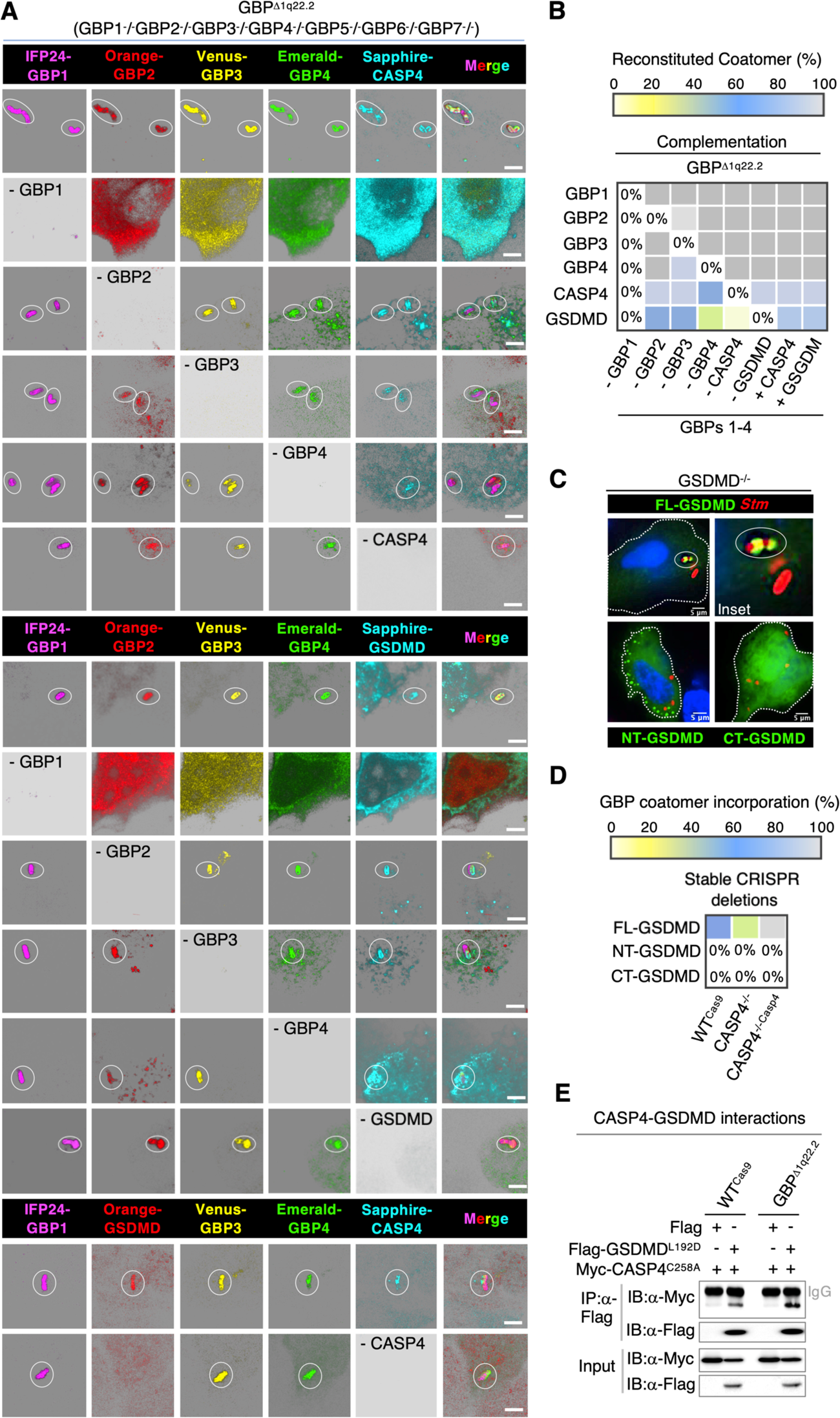
GBP1-dependence for 6-member coatomer assembly and caspase-4 recruitment of GSDMD. (**A**) Multicolor confocal imaging of the reconstituted coatomer in the septuple GBP*^Δ^*^1q22.2^ mutant. Host cells were not activated to avoid endogenous IFN-*γ* induced caspase-4 expression affecting assembly in the top and bottom panels. Stepwise omission of each coatomer component revealed clear dependence on GBP1. For the bottom panel, Orange-GBP2 was exchanged with Orange-GSDM since GBP2 is dispensable for reconstituted coatomer formation. Scale bar, 2μm. (**B**) Heat map of *Stm* recruitment by individual coatomer components in the presence of GBPs 1-4. Left to right, depicts the individual component subtracted from or added to the GBP1-4 core module. (**C**) Widefield imaging of full-length (FL), N-terminal or C-terminal fragments of human GSDMD reintroduced into GSDMD^-^/^-^ cells activated with IFN-*γ* and infected with *Stm* for 2 hours. Clear targeting of full-length GSDMD to bacteria was evident (inset, dashed circle). Nuclear staining with DAPI. Scale bar, 5 (**D**) GSDMD targeting was largely reliant on caspase-4 *in situ* as shown in IFN-*γ*-primed CASP4^-^/^-^ cells with or without complementation. Scale bar, 5μm. (**E**) Co-immunoprecipitation of Flag-GSDMD^L192D^ by Myc-CASP4^C258A^ (to reduce cytotoxicity; 28) in both wild-type HeLa CCL2 cells as well as in the septuple GBP*^Δ^*^1q22.2^ mutant. IgG, immunoglobulin heavy chain.

**Fig. S4.**
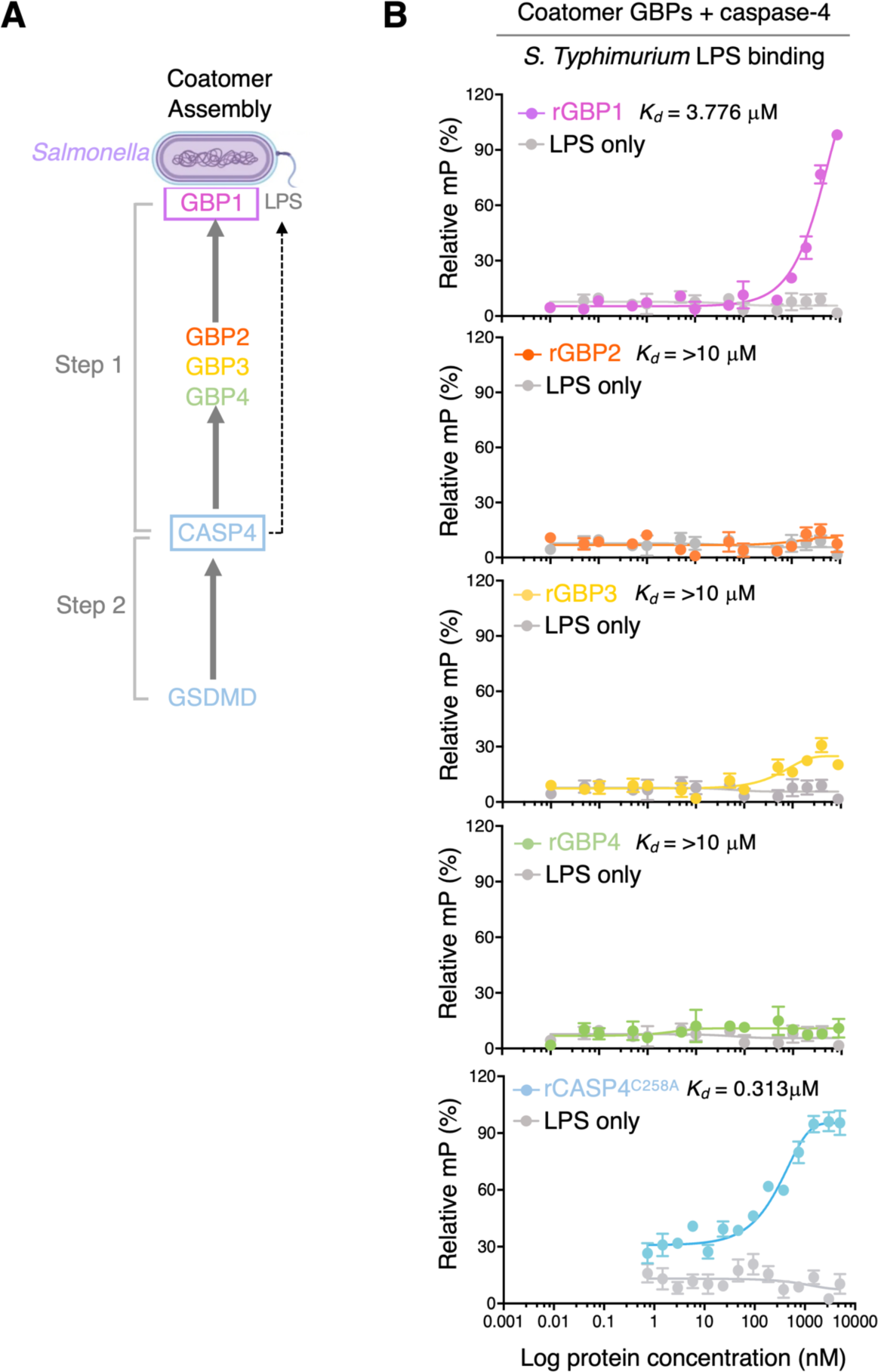
Complete 6-member coatomer assembly with GBP1 and caspase-4 the primary LPS-binding proteins. (**A**) Two step model of coatomer assembly where GBP1 and caspase-4 control initially GBP2-4 and subsequent GSDMD recruitment, respectively. Pro-IL-18 recruitment to the coatomer has infrequently observed (< 1-3%; data not shown) and hence questionable that it is a *bone fide* coatomer member. (**B**) *Salmonella* minnesota LPS-Alexa Fluor 488 (ThermoFisher) binding curves for recombinant coatomer proteins in fluorescence anisotropy assays. Y-axis, % of maximal polarization (mP). Mean ± SD determined in triplicate for each protein concentration. Representative of 3 independent experiments.

**Fig. S5.**
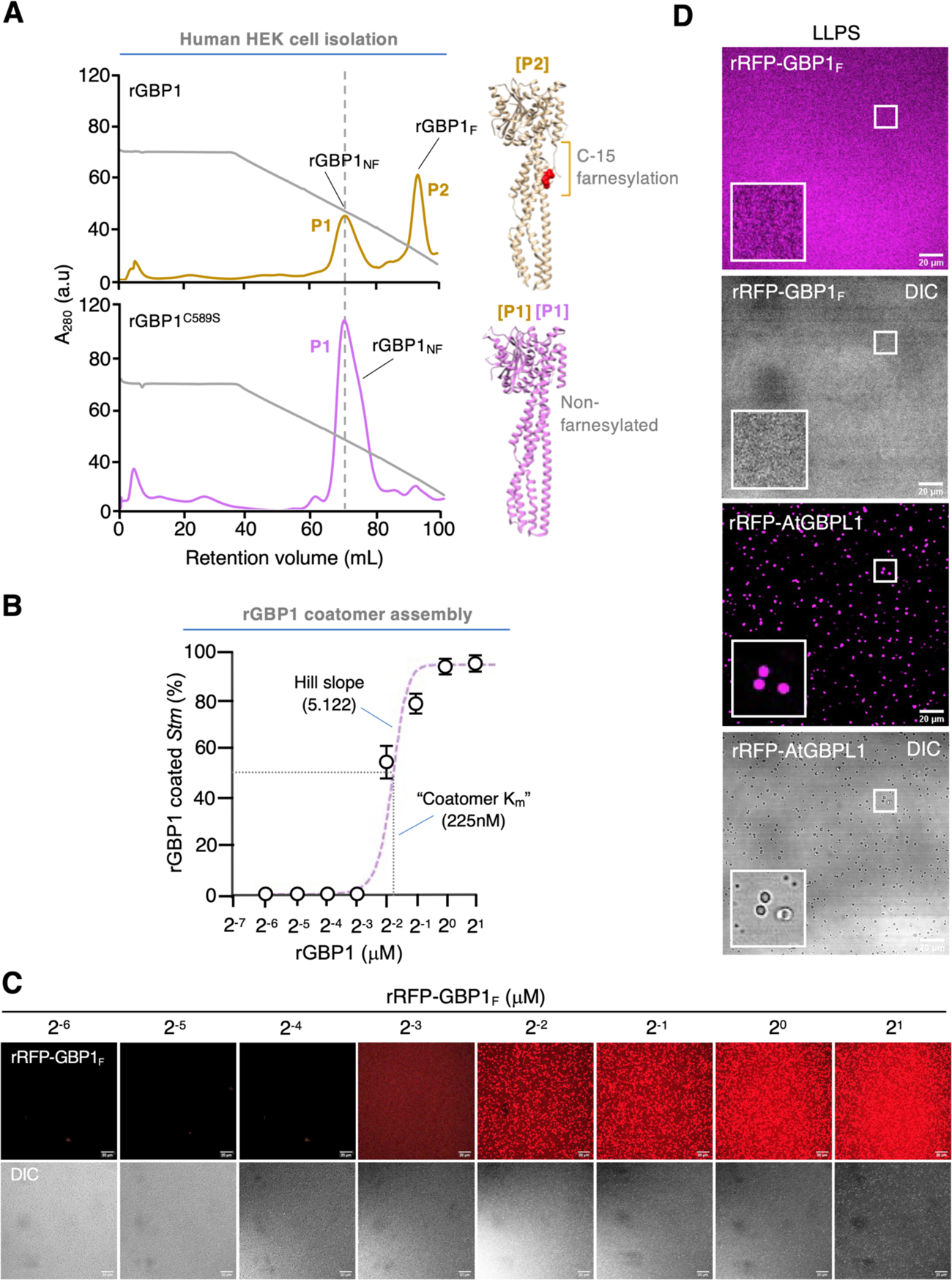
Reconstituted assembly characteristics with farnesylated recombinant GBP1 purified from human cells. (**A**) Isolation of lipidated human RFP-GBP1 for coatomer studies. Superdex-600 FPLC preparative profiles of isolated peaks containing farnesylated (peak 2; P2) and non-farnesylated (peak 1; P1) species; recombinant RFP-GBP1^C589S^ served as an internal non-farnesylated control. Right, position of the C-15 farnesyl moiety at a final C-terminal branch in the 1F5N crystal structure. (**B, C**) rRFP-GBP1 coatomer assembly measured on heat-inactivated *Stm* 1344 to ensure immobility during the assay. 2 *µ*M rRFP-hGBP1 plus 2 mM GTP tested across increasing diluents (below) imaged by confocal microscopy. Best-fit interpolation curve fitting of means ± SD together with regression analysis found half maximal (population-based “coatomer K_m_”) and Hill slope values using Graph Pad Prism 9.1.1. (**D**) Liquid-liquid phase separation (LLPS) droplet assay without Ficoll for rRFP-GBP1 and rRFP-AtGBPL1, the latter as a positive control. Pseudo-colored scanning confocal images collected with a 63×/1.40 oil immersion objective. Inset, coacervated droplets evident for rRFP-AtGBPL1 but not rRFP-GBP1. One of two representative experiments shown.

**Fig. S6.**
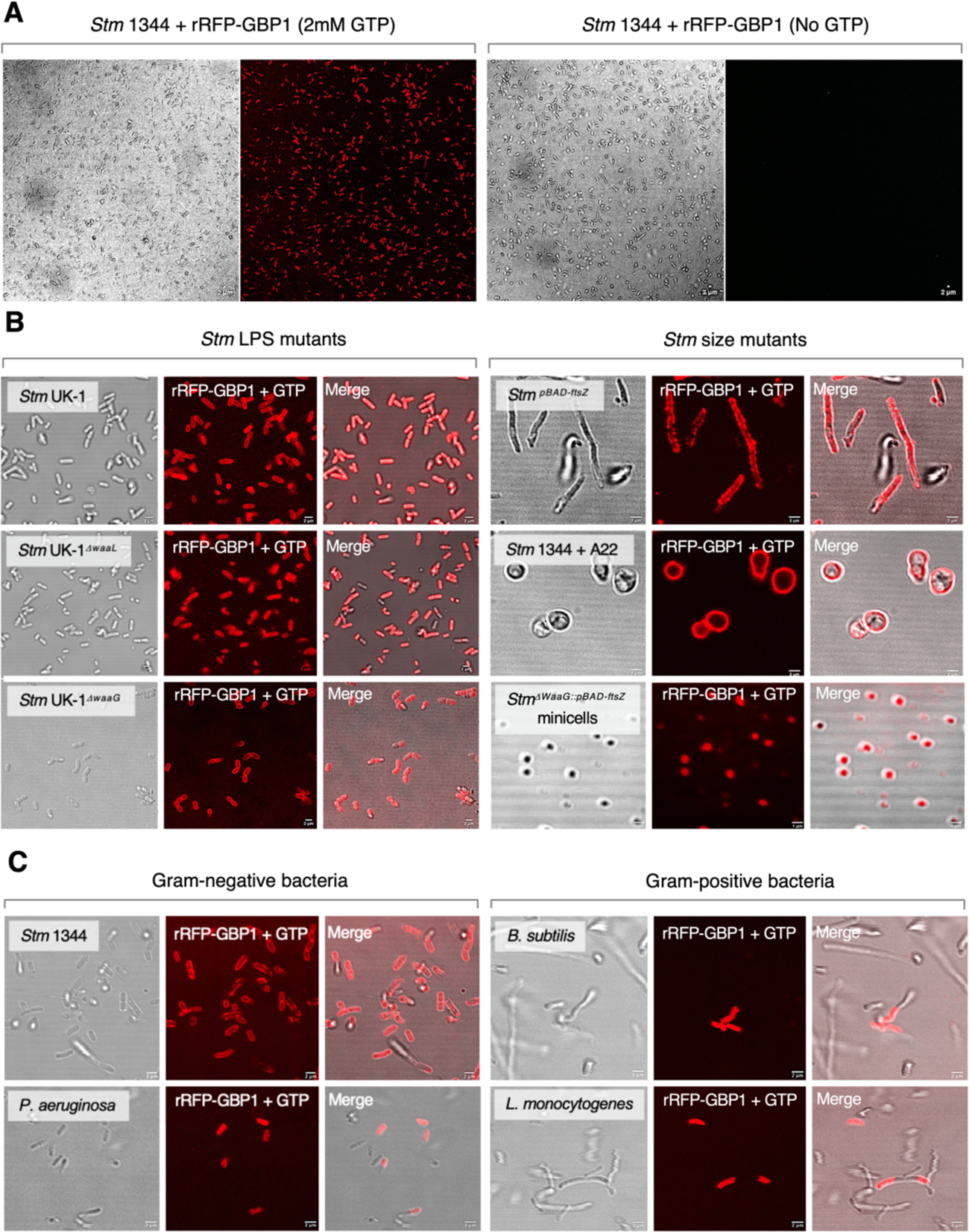
Reconstituted coatomer on different bacterial mutants and species. (**A**) GTP-dependent reconstituted coating of rRFP-GBP1 on *Stm* 1344 *in vitro*. Scanning confocal images using 63×/1.40 oil immersion objective. No assembly observed without GTP. Scale bar, 2μm. (**B**) GTP-dependent rRFP-GBP1 coatomer reconstitution using *Stm* UK-1 LPS mutants (left) or size variants (right). The latter included genetically altered *Stm^pBAD::ftsZ^* elongated strains, A22-treated (S-(3,4-dichlorobenzyl)isothiourea; 500μM) circular strains, and purified *Stm^ΔWaaG::pBAD-FtsZ^* minicells. Scale bar, 2μm except minicells (1μm). (C) Coatomer reconstitution on Gram-negative and Gram-positive bacterial strains. Scale bar, 5μm.

